# Influence of Geographical Aspect and Topography on Canopy Openness in Tropical Rainforests of Sri Lanka along an Altitudinal Gradient

**DOI:** 10.1101/2023.07.06.547768

**Authors:** R.M.C. Madhumali, W.M.P.S.B. Wahala, H.K.N. Sanjeewani, D.P. Samarasinghe, W.A.J.M. De Costa

## Abstract

Solar radiation energy is a limiting factor for forest growth in humid tropical environments. Radiation incident on a forest canopy varies with azimuth angle of incident radiation and aspect of the forest terrain. The objective of this work was to determine the influence of the geographical aspect and azimuth of incident radiation on the canopy structure of tropical rainforests of Sri Lanka (TRFSL). Hemispherical photography was used to capture canopy images of TRFSLs in ten permanent sampling plots across an altitudinal range from 117 m to 2132 m above mean sea level. Fraction of visible sky (V_sky_) in 144 sectors of the canopy hemisphere, defined by 18 zenith angle (ZnA) × 8 azimuth angle (AzA) segments, was determined using image analysis. Canopy openness, quantified as V_sky_ of the overall hemisphere, increased with increasing altitude. Canopy leaf area index, decreased linearly with altitude and had a negative relationship with V_sky_. Canopy openness of the top one-third (ZnA=0°–30°) of the hemisphere (V_sky(0-30)_) was significantly (*p*<0.05) lower on the east (AzA=90°) than on the west (AzA=270°) in a majority of plots. Similarly, V_sky(0-30)_ was lower on the northern (AzA=0°) than the southern (AzA =180°) canopy segment. These patterns were altered in plots where nearby mountains and slope aspect of the plot influenced incident radiation. These observations suggest a hypothesis that canopies of TRFSL are structured to maximize radiation capture by allocating more leaf area, and therefore having lower canopy openness, on sides of the canopy which face directions of greater radiation receipt.

## INTRODUCTION

Capture of solar radiation by a forest canopy is essential for photosynthesis, biomass production and forest growth. Canopy openness, defined as the unobstructed visible sky fraction through a forest canopy when viewed from the ground level, determines the light environment within and below a forest canopy (Chazdon and Fetcher, 1984; Gonsamo *et al*., 2013). Canopy openness also influences forest regeneration in canopy gaps and thereby determines the species composition and complexity of the forest structure (Denslow, 1987; Yamamoto, 1992; Parker, 2004; Marler and del Moral, 2018; Mazon *et al*., 2020). The amount of light that penetrates the forest canopy is determined by the extent of canopy openness and the rate of gap creation, which influences the microclimate within the forest (Jennings *et al*., 1999; McCarthy and Robison, 2003). Seed germination and sapling growth depend upon light that reaches the forest floor, which acts as a limiting resource for plant growth (Oberbauer *et al*., 1988; Goodale *et al*., 2012; Unger *et al*., 2013).

Topographical measures are important when studying canopy openness and solar radiation regimes of forest canopies. Radiation energy load incident on a forest canopy varies with latitude, aspect (i.e. compass direction) of the terrain and its slope (Holland and Steyn, 1975; Liang *et al*., 2006; Gallardo-Cruz *et al*., 2009). Penetration of incident radiation in to the forest canopy and its distribution within the forest is determined by the canopy attributes such as leaf area index, morphology and architecture (McCarthy and Robinson, 2003; Parker, 2014). Leaf area index is defined as one-sided leaf area per unit ground area and it acts as the exchange surface for photosynthesis and transpiration (Breda, 2003; Unger, 2013). It drives canopy carbon production and carbon sequestration, water balance of the site, biogeochemical cycles and the microclimate within the forest (Breda, 2003; Woodgate *et al*., 2015).

Different levels of radiation receipt on different aspects of a slope have been shown to induce appreciable variation in the forest ecosystems on opposing sides. For example, Smith (1977) observed higher solar irradiance and higher maximum temperatures on east-facing slopes than west-facing slopes in the grasslands of Mount Wilhelm of Papua New Guinea because of clear mornings and cloudy afternoons. Smith (1977) further observed significant differences in species composition between the east- and west-facing slopes. Smith *et al*. (1992) observed that canopy openness in the eastern and western sectors of canopy hemispherical images of a mature lowland moist tropical forest in Panama were not correlated. This indicated the possibility of the existence of different microsites with different radiation regimes on different aspects of a forest canopy. Solar radiation and temperature regimes have also been reported to be different on north- and south-facing slopes of mountains and ridges (e.g. Cantlon, 1953), which have been linked to changes in vegetation. For example, Marler and del Moral (2018) reported appreciable differences in vegetation composition and structure on slopes facing different aspects in the tropical landscapes below the Mount Pinatubo in the Philippines. Holland and Steyn (1975) reported greater extent of montane forests on wetter southern slopes than drier northern slopes of tropical East African mountains. Liang *et al*. (2006) reported lower growth rates of selected tree species on the western-than eastern slopes of the north-east Tibetan Plateau. Holland and Steyn (1975) provide several examples of environmental variations with slope aspect which have resulted in clear differences in vegetation. On the other hand, Gallardo-Cruz *et al*. (2009) observed that the vegetation structure and β-diversity of a seasonally-dry tropical landscape in Mexico were not related to environmental changes associated with slope aspect and altitude. However, there is a paucity of studies which have investigated the influence of aspect on vegetation properties, especially the canopy properties, in tropical forests.

Most tropical rainforests of Sri Lanka (TRFSL) are located on the south-western slopes and the central highlands with substantial within-forest variation in slope aspect coupled with highly heterogeneous topography. However, there has been no previous work that has investigated the influence of geographical aspect on canopy properties such as leaf area index, openness and structure in Sri Lankan rainforests. Therefore, the principal objective of the present work was to determine the possible influence of geographical aspect and azimuth of incident radiation on the canopy structure of TRFSL.

## METHODOLOGY

### Study Sites

The study was done in ten permanent sampling plots (PSPs) of 1 ha (100 m × 100 m) extent in selected tropical rainforests in the wet zone of Sri Lanka along an altitudinal gradient from 117 m to 2132 m above mean sea level (amsl). Two plots each were located in the forest reserves at Kanneliya (KDN 1 and KDN 2 at 117 and 174 m *amsl*), Sinharaja-Pitadeniya (PTD 1 and PTD 2 at 509 and 618 m) and Sinharaja-Enasalwatte (ENS 1 and ENS 2 at 1042 and 1065 m) (Figure 1). One PSP each was in the forest reserves at Rilagala (RLG at 1668 m), Hakgala (HKG at 1804 m), Pidurutalagala (PTG at 2080 m) and Horton Plains (HTN at 2132 m). Long-term climatic data of the 10 PSPs are given in Madhumali *et al*. (2021). Soils of all plots have derived from Igneous and Gneissic rocks and belong to the Red Yellow Podzolic group according to the local soil classification (Panabokke, 1996; Mapa *et al*., 1999). According to the USDA soil taxonomic classification, soils of the PSPs belong to Typic Plinthudults (KDN 1 and 2), Typic Haplohumults (PTD 1 and 2, ENS 1 and 2), Typic Hapludults (RLG), Typic Paleudults (HKG and PTG) and Typic Humitropents (HTN) (Mapa et al., 1999). According to the FAO/UNESCO soil classification, the soils belong to Dystric Plinthosols (KDN 1 and 2), Humic Alisols (PTD 1 and 2, ENS 1 and 2, RLG, HKG and PTG) and Dystric Cambisols (HTN) (Mapa et al., 1999). Topographic details of plots (i.e. aspect and slope) are given in Table 1. Forests in all PSPs except those at Kanneliya were at the climax or late-successional stage and free from anthropogenic disturbance and dieback during the last 40 years (Madhumali *et al*., 2021). The Kanneliya Forest Reserve had been selectively logged from 1968 to 1988 (Gunawardena, 2019) and was in the middle of secondary succession. However, Kanneliya was included in the study because of its high floristic diversity and low altitude, which enabled a wide altitudinal range for the study.

**Figure 1:**
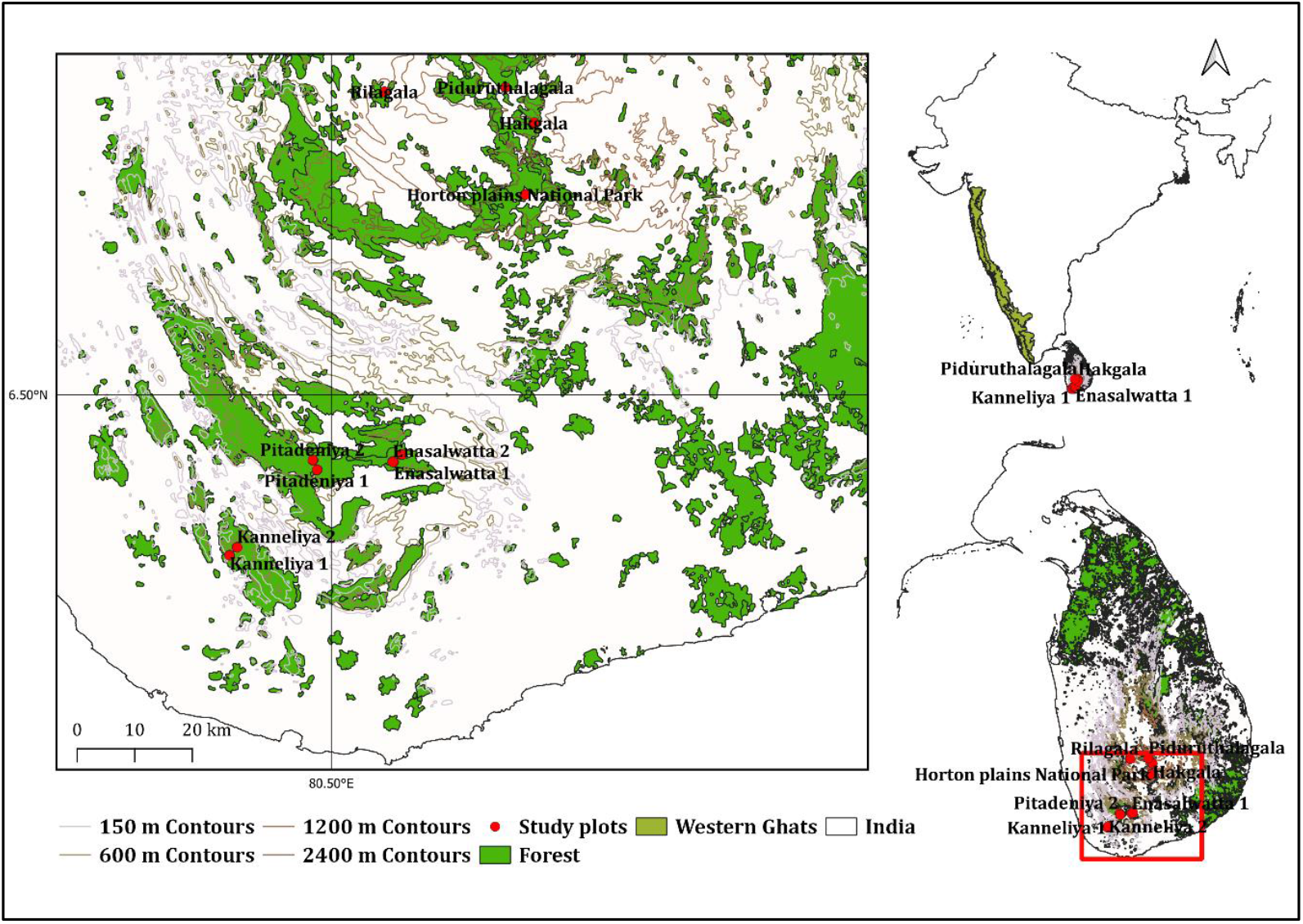
Locations of permanent sampling plots in tropical rainforests of Sri Lanka along an altitudinal gradient from 117 m to 2132 m above mean sea level. See Table 1 for further description of study sites and sampling plots.

**Table 1:** Site description along with their topography and aspect

| PSP<sup>†</sup> | Altitude<br>(m *amsl*) | Aspect | Slope<br>(%) | Obstructions to incident radiation |
| --- | --- | --- | --- | --- |
| KDN 1 | 117 | West-facing | 16 | Mountain of 300 m height<sup>‡</sup> on the East at 400 m away |
| KDN 2 | 174 | North-west facing | 7 | Mountain on the East (100 m) at 500 m away and on the West (150 m) at 300 m away |
| PTD 1 | 509 | East-facing | 13 | Mountain on the East (100 m) at 500 m away and mountain on the West (140 m) within 300 m away |
| PTD 2 | 618 | West-facing | 9 | Mountain on the East (120 m) and on the West (50 m) within 400 m |
| ENS 1 | 1042 | East-facing,<br>South-facing | 11,7 | No obstructions nearby |
| ENS 2 | 1065 | East-facing,<br>North-facing | 14, 5 | No obstructions nearby |
| RLG | 1668 | East-facing | 15 | No obstructions nearby |
| HKG | 1804 | North-east facing,<br>North-facing | 22,14 | Mountain on the South-west (217 m) at 300 m away |
| PTG | 2080 | West-facing | 19 | Mountain on the East (100 m) 100 m away |
| HTN | 2132 | West-facing | 9 | No obstructions nearby |

<sup>†</sup>PSP – Permanent Sampling Plots; KDN 1, KDN 2 – Kanneliya Forest Reserve Plot 1 and 2; PTD 1, PTD 2 – Sinharaja-Pitadeniya Plot 1 and Plot 2; ENS 1, ENS 2 - Sinharaja-Enasalwatta Plot 1 and Plot 2; RLG - Rilagala Forest Reserve; HKG – Hakgala Strict Nature Reserve; PTG – Pidurutalagala Forest Reserve; HTN – Horton Plains National Park. <sup>‡</sup>Height of the mountain has been estimated relative to the elevation of the PSP.

### Canopy Hemispherical Photography (CHP): Image Acquisition and Processing

Each 1 ha plot was demarcated in to 25 m × 25 m sub-plots by a grid of marked poles producing nine sampling points to obtain hemispherical photographs of the forest canopy. A hemispherical photograph of the forest canopy gives an upward-looking image of the sky through the forest canopy with a 180° field of view. As such it provides a permanent record of the sky geometry of visibility and obstruction at the time of image acquisition (Anon, 1999). Therefore, CHP provides a non-destructive and non-invasive method to calculate forest canopy parameters and solar radiation regimes within the forest. Canon 80D digital SLR camera with a Sigma EX DC 4.5 mm F2.8 circular fisheye lens was fixed to a self-levelling mount with a tripod. This setup was placed 1.3 m above the ground on each sampling point, aligned to north when the image was taken at each of the nine sampling points in each PSP. Focal length was set to infinity and images were taken under auto exposure settings. Further details of the camera settings can be found in Madhumali *et al*. (2021). Most of the time images were taken in the morning to minimize bright sun patches. Three full rounds of Images, each round covering all 10 PSPs and consisting of 90 images, were taken during a 16-month period from September 2019 to January 2021. The interval between two consecutive rounds varied from 3 to 5 months, depending on the location.

Canopy hemispherical images were processed using the HemiView Canopy Analysis Software (Version 2.1, Delta-T Devices, Cambridge, UK). Prior to analysis, non-canopy objects of the image (stems and soil) were excluded using Adobe Photoshop (version 11.0). Lens properties and solar model properties were also set before the analysis of images. A detailed description of these properties can be found in Madhumali *et al*. (2021). The magnetic north of the image was aligned and the particular date and study site was selected for each specific image. The images were classified based on a threshold value. The optimum threshold value gives dark parts as canopy material (i.e. mainly leaves, twigs and smaller branches) and white parts as the sky. Every image was analysed three times by the same person to obtain a precise result.

### Calculation of Canopy Properties

In a canopy hemispherical image, HemiView demarcates a sky sector by a 5° (zenith angle) × 45° (azimuth angle) combination (Figure 2). Accordingly, HemiView defines 144 sky sectors as combinations of 18 zenith angles × 8 azimuth angles. Visible sky fraction for a given sky sector identified by a zenith angle, θ, and an azimuth angle, α, (V_sky(θ,α)_) is defined as the proportion of visible sky fraction in the sky sector, with respect to the entire hemisphere (Anon, 1999).

**Figure 2:**
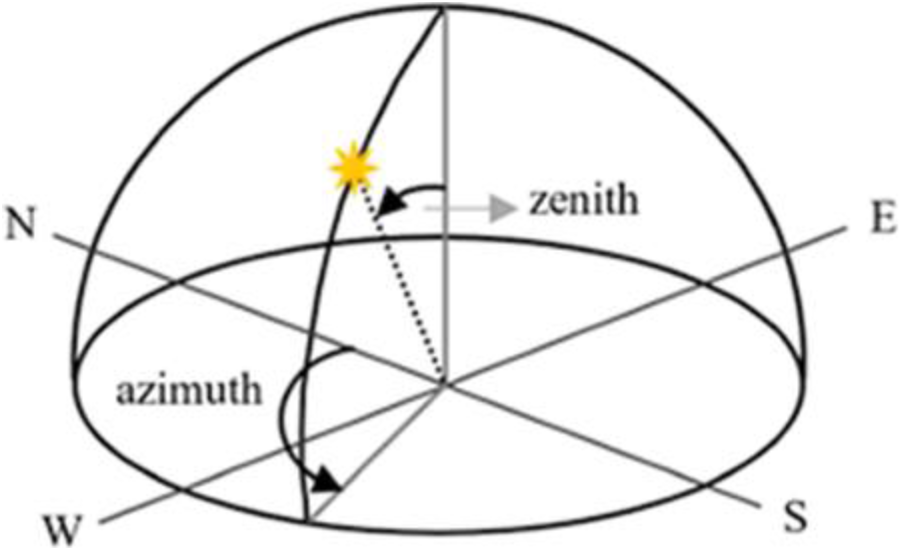
Zenith and Azimuth angles of the canopy hemisphere.

Visible sky fraction of the entire canopy image (V_sky_) is calculated as,

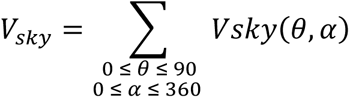

Leaf area index of each sky sector (LAI_(θ,α)_) is computed by inverting the Beer-Lambert law of radiation penetration as,

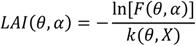

where, F is the sky gap fraction of each sky sector, k the light extinction coefficient and X the ellipsoidal leaf angle distribution parameter (Anon, 1999; Madhumali *et al*., 2021). Gap fraction of a given sky sector is the fraction of visible sky within the sky sector. Summation of LAI_(θ,α)_ values of all sky sectors are calculated as LAI of the whole canopy.

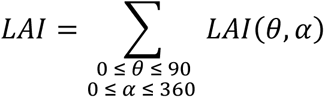

V_sky_ was taken as an overall measurement of the ‘openness’ of the entire forest canopy. Contour maps of canopy openness were developed for the entire canopy by mapping the variation of V_sky(θ,α)_ across the 144 sky sectors. For developing the contour maps, mean V_sky(θ,α)_ for each sky sector was calculated by averaging the respective values in all three measurement rounds and all nine sampling points in each PSP. Canopy openness of the four major aspects (i.e. North, East, South and West) were extracted by selecting the respective sets of V_sky(θ,α)_ values at azimuth angles of 0°, 90°, 180° and 270° respectively. Canopy openness values were also extracted for three zenith angle segments as 0°-30°, 30°-60° and 60°-90°. The interpretation of these three zenith angle segments is given along with their results in the Results and Discussion section.

### Estimation and measurement of the differential irradiance and shading on the permanent sampling plots during pre- and post-noon periods

Differential irradiance and shading of forests in the PSPs during the periods from sunrise to solar noon (pre-noon period) and from solar noon to sunset (post-noon period) were mapped using Digital Elevation Model (DEM) images from SRTM 1 Arc-Second Global which were downloaded from the USGS Earth Explorer website (https://earthexplorer.usgs.gov/). Resolution of the image was 1 arc second (approximately 30 m). Images acquisitioned on 2014.09.23 were used. The DEM image was imported to Arc map version 10.8 and ‘hill shade’ analysis was done to map pre-noon and post-noon shadows on each of the ten PSPs. ‘Hill shade’ creates shaded relief from a surface raster by considering the source illumination angle which is the sun position as defined by the azimuth angle (0°-360°) and altitude angle above the horizon (0°-90°). Hill shade analysis was studied at sun angles of 20°, 30° and 45° for morning and evening.

Automated weather stations (AWSs) had been established in open areas near the PSPs at Kanneliya (closer to KDN 1 than to KDN 2), Sinharaja-Enasalwatte (approximately equal distance from ENS 1 and ENS 2) and Pidurutalagala. A pyranometer on the AWSs recorded the incident total solar irradiance at one-minute time intervals. Diurnal variation of instantaneous solar irradiance on selected days of high daily total solar irradiance was examined to determine the differential irradiance levels during the pre- and post-noon periods.

### Statistical Analysis

#### Variation of canopy openness (V_sky_) and leaf area index (LAI) among permanent sampling plots (PSPs) and measurement rounds

Significance of the variation of canopy openness (V_sky_) and LAI among different PSPs and measurement rounds were tested by analysis of variance (ANOVA) in a general linear model using Proc GLM in SAS^®^ Studio (SAS Institute Inc., 2021). Pooled data set of each variable was tested for normality using the Shapiro-Wilks test statistic (*W*) in Proc Univariate of SAS. When the data were not normally-distributed, they were transformed using the optimum power transformation found by the Box-Cox method (Box and Cox, 1964).

#### Variation patterns of V_sky_ and LAI with altitude

Variation of V_sky_ and LAI with altitude was determined by linear and polynomial regression. Initially, regressions were fitted for different measurement rounds separately. When these regressions did not show differences among rounds, data from all rounds were pooled and regressions were done.

#### Relationship between V_sky_ and LAI

The relationship between V_sky_ and LAI was determined by fitting of a power function of the form, V_sky_ = a (LAI)^b^. The parameters *a* and *b* were determined by doing a linear regression between log_10_(V_sky_) and log_10_(LAI).

#### Variation of canopy openness along different zenith angle segments and selected azimuth angles of the canopy hemisphere

Significance of the variation of V_sky(θ,α)_ values of the three zenith angle segments (0°-30°, 30°-60° and 60°-90°) and the azimuth angles representing the four major aspects (i.e. North, East, South and West) was determined by a linear mixed model (LMM) with repeated measures analysis with Proc Mixed in SAS using the residual restricted maximum likelihood (REML) method for estimating the covariance structure. As this study focused on determining the variation of V_sky_ across the whole range of altitudes rather than at the specific altitudes of the PSPs, altitude (β) was considered as a random effect. As such, the altitudes of PSPs were considered as random points along the range of altitudes occupied by tropical rainforests of Sri Lanka. On the other hand, the three zenith angle segments (θ) and the four azimuth angles (α) representing the four directions had been purposively selected. As such, θ and α were considered as fixed effects in the LMM analysis. As the measurements of V_sky_ on all PSPs and their fixed sampling points were repeated over three rounds, the measurement round (γ) was considered as a repeating factor. The covariance structure of the random effect was specified as the variance component (VC) type. To determine the influence of θ and α in each PSP, the LMM analysis was performed separately for each PSP. Least square (LS) means of V_sky(θ,α)_ and their confidence intervals (CIs) were computed for determining the significance of the effects of α (i.e. aspect) in different segments of θ (i.e. zenith angle segments). When the CIs of two LS means did not overlap, the two LS means were considered significantly different.

## RESULTS AND DISCUSSION

### Variation of Canopy Openness (V_sky_) and Leaf Area Index (LAI) among Permanent Sampling Plots (PSPs) and Measurement Rounds

Normality testing of the data sets of V_sky_ and LAI pooled across the three measurement rounds, 10 PSPs and nine sampling points (Supplementary Figure S1.a and b) showed that they deviated significantly from normality (V_sky_: Shapiro-Wilks *W* = 0.953, *p*<0.0001, n = 270; LAI: Shapiro-Wilks *W* = 0.794, *p*<0.0001, n = 270) (Supplementary Figure S1). The distribution of LAI data deviated significantly from normality even after removing one clear outlier (i.e. 8.90) also (Shapiro-Wilks *W* = 0.887, *p*<0.0001, n = 269). Data sets were normalized by Box-Cox power transformations of λ = 0.25 for V_sky_ and λ = -1 for LAI.

Analysis of variance of transformed data revealed significant PSP × measurement round interaction effects on both V_sky_ (*p*<0.0001) and LAI (*p*=0.0025). Both V_sky_ and LAI showed highly-significant (*p*<0.0001) variation among PSPs. On the other hand, variation among different rounds of measurement was significant only for LAI (*p*=0.0191), but not for V_sky_ (*p*>0.05). Significance of main effects (*p*<0.0001 and *p*=0.0306 for PSP and round respectively) and the interaction effect (*p*=0.0036) did not change for LAI data when ANOVA was done after removing the outlier. In all ANOVAs, neither V_sky_ nor LAI differed significantly (*p*>0.05) among different sampling points within a plot.

Plot-level means of V_sky_ and LAI at a given altitude did not differ significantly (*p*>0.05) between different measurement rounds at a majority of altitudes (Figure 3). This showed that canopy openness and size did not vary during the 16-month period of the present study. The absence of prolonged rain free periods in the humid, tropical climatic zone (Woodward, 1987) in which all forest plots are located is responsible for the absence of seasonal variation in canopy size and structure. This is in accordance with Borchert (1988). As all plots, including the two at Kanneliya, are within mature, late-successional forests, it is highly likely their canopies have already reached or are close to the upper-limit of canopy size (i.e. LAI) that their respective climates and soil resource availability levels permit. In particular, the forest canopies in plots from lower to mid altitudes from Kanneliya to Sinharaja-Enasalwatte have completed canopy closure and vertical structuring, which is typical for tropical wet evergreen rainforests (Oldeman, 1983; Torquebiau, 1986; Gunatilleke and Ashton, 1987). The range of LAI values observed in the present work were within the global-scale ranges reported for tropical rainforests (Asner, 2003; Iio *et al*., 2014) and tropical montane forests (Moser *et al*., 2007; Heiskanen *et al*., 2015).

**Figure 3:**
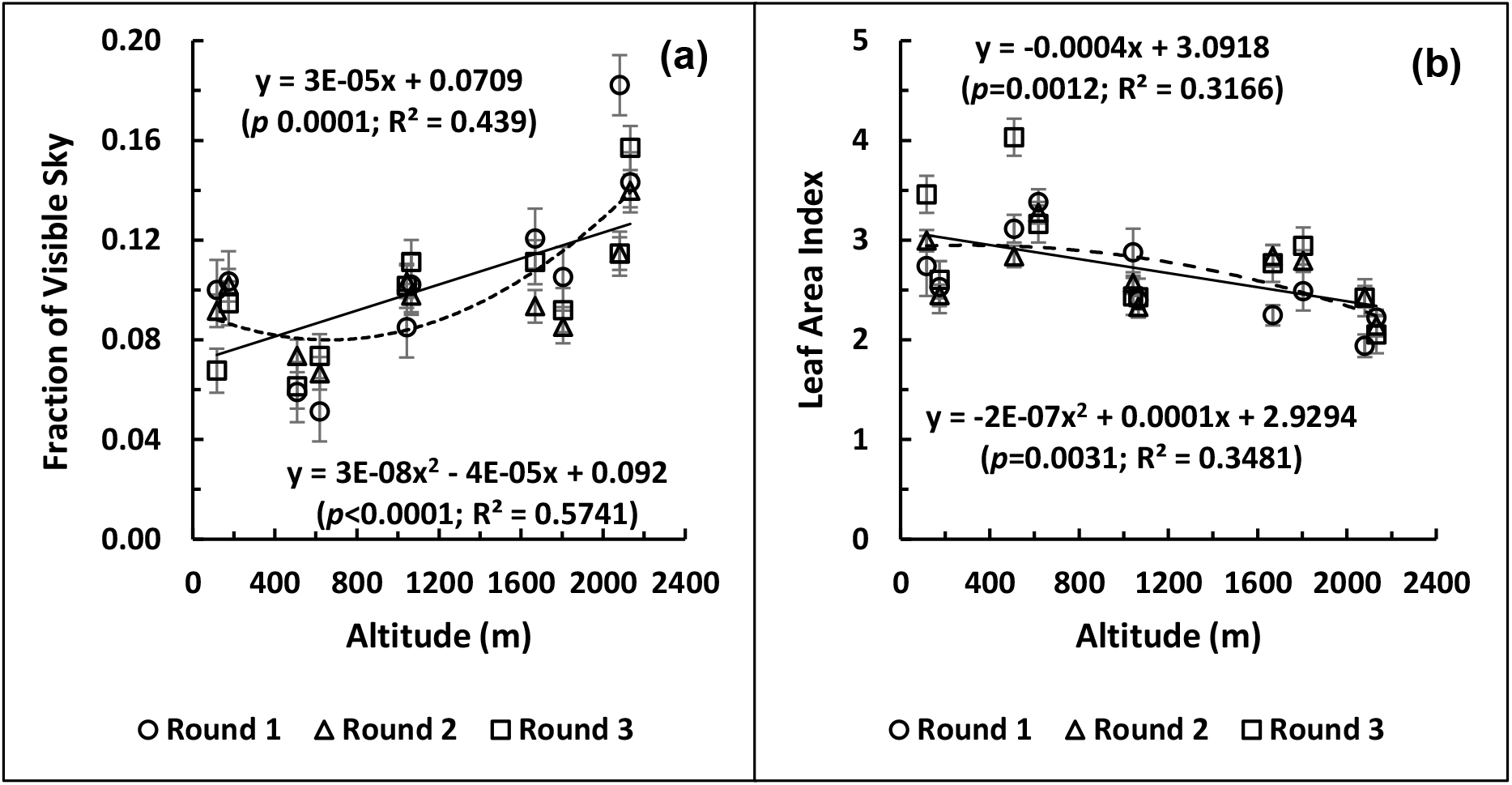
Variation of canopy openness (Fraction of Visible Sky, V_sky_) (a) and leaf area index (LAI) (b) with altitude. Linear and second-order polynomial functions are shown by solid and broken lines respectively.

### Variation Patterns of V_sky_ and LAI with Altitude

Mean V_sky_ values of Round 3 at 116 m, Round 2 at 1668 m and Round 3 at 2032 m were the only ones which differed significantly from the respective means of the other two rounds (Figure 3.a). The corresponding mean LAI values which showed significant variation from the rest at a given altitude occurred in Round 1 at 1042, 1668 and 2080 m and in Round 3 at 116 and 174 m (Figure 3.b). Accordingly, common regression models were fitted after pooling the respective data from all three rounds to determine the response patterns of V_sky_ and LAI to altitude. Linear functions showed highly-significant fits for the increase of V_sky_ (*p*<0.0001; *R^2^* = 0.439) and for the decrease of LAI (*p*=0.0012; *R^2^* = 0.317) with increasing altitude. Second-order polynomial functions, with minimum V_sky_ at 663 m *amsl* and maximum LAI at 302 m, appreciably improved the accuracy of fit in the regression of V_sky_ (*p*<0.0001; *R^2^* = 0.574), whereas only a marginal improvement could be observed for LAI (*p*=0.0031; *R^2^* = 0.348). Similarly, excluding the outlier in the LAI data set only marginally improved the accuracy of fits with the linear (*p* = 0.0006; *R^2^* = 0.347) and polynomial (*p* = 0.0016; *R^2^* = 0.380; maximum LAI at 285 m *amsl*) regression models (Supplementary Figure S2). The linear decreasing trend of LAI across the whole altitudinal range is in agreement with decreasing trends observed by Moser *et al*. (2007), Unger *et al*. (2013) and Pfeifer *et al*. (2012) for tropical forests in Africa and South America. In a detailed analysis of the influence of long-term climatic variables, Madhumali *et al*. (2021) showed that decreasing annual total rainfall across the altitudinal gradient of the PSPs had a dominant influence on the decreasing trend of LAI, with decreasing mean annual air temperature and annual total solar irradiance also having less strong effects. This is in agreement with the conclusions of the global-scale analysis of Iio *et al*. (2014).

Photosynthetic rates at higher altitude forests are constrained by the lower temperatures and the poor capacity of high-altitude tropical trees to acclimate to increasing temperatures (Vårhammar et al., 2015; Dusenge et al., 2021; Wittemann et al., 2022). Therefore, the pool of primary assimilates from which a portion is allocated for leaf production would be smaller. Furthermore, leaf thickness, as quantified by leaf mass per unit area (LMA), increases with increasing altitude (van de Weg et al, 2009; Asner and Martin, 2016). This indicates that leaf construction costs are higher in high-altitude TRFs (i.e. montane forests). Therefore, despite the lower maintenance costs at lower temperatures, the decreasing carbon pool, the consequent decrease in the carbon allocation for leaf growth and increasing leaf construction costs could be responsible for the observed reduction in LAI with altitude. Additional stress factors that are encountered in the high-altitude montane forests exert further limitations on leaf growth and LAI in tropical montane forests. For example, high wind speeds and the consequent high transpiration rates and water stress require adaptations such as smaller and thicker leaves with stronger cell walls, thus restricting the LAI of such forest canopies.

### Relationship between V_sky_ and LAI

When the data from all rounds, PSPs and sampling points were pooled, there was a negative curvilinear relationship between V_sky_ and LAI (Figure 4).

**Figure 4:**
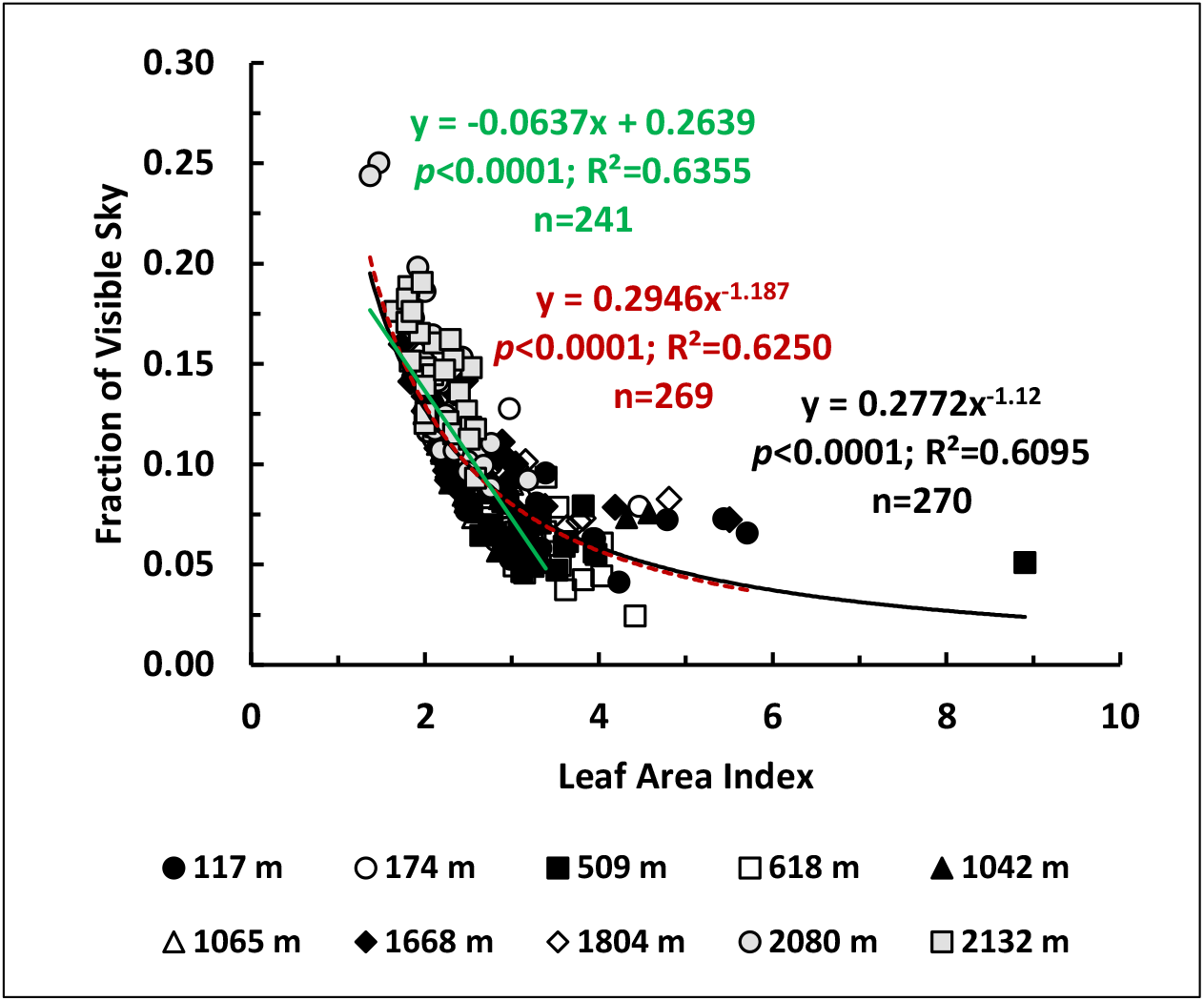
Relationship between Fraction of Visible Sky (V_sky_) and Leaf Area Index, (LAI). Non-linear functions are fitted with-(solid, black curve) and without (broken, maroon curve) the outlier (LAI>8). Linear function is fitted for LAI ≤ 3.5. Data points represent individual canopy hemispherical images in three measurement rounds over a 16-month period at nine sampling points each in ten permanent sampling plots across a range of altitudes.

Canopy openness, as quantified by V_sky_, decreased with canopy size, almost linearly at a rate of 0.06 per unit increase of LAI up to a value of around 3.5. At higher LAIs, a very slow decline of V_sky_ could be observed. The increase in slope with decreasing LAI (i.e.greater increase in V_sky_ per unit decrease in LAI) shows the canopy opens faster as its size decreases in forests at higher altitudes.

### Distribution of Canopy Openness along Zenith and Azimuth Angles of the Canopy Hemisphere

Contour maps showing the distribution of canopy openness in different sky sectors defined by zenith (θ) and azimuth (α) angle segments (V_sky(θ,α)_) of the canopy hemisphere are shown in Figure 5 for the 10 PSPs. Contours of V_sky(θ,α)_ between zenith angles from 0° to 30°, which represent the upper one-third of the hemisphere, and therefore, the canopy gaps through which radiation penetrates during the middle period of the day (i.e. 2-3 hours around noon), showed an increasing trend from the lower altitudes (i.e. Kanneliya, 117 – 174 m, and Sinharaja-Pitadeniya, 509 – 618 m) to higher altitudes (i.e. Pidurutalagala and Horton Plains, 2080 – 2132 m). Out of the four lower altitude plots at Kanneliya and Pitadeniya, KDN 1 at 117 m showed a greater openness at the upper one-third of the hemisphere than the others. The middle (zenith angle from 30° to 60°) and lower (60° to 90°) thirds of the hemisphere showed progressively lower canopy openness because of the accumulation of leaf layers and stems in the hemispherical photograph (Supplementary Figure S3). This reduction in V_sky(θ,α)_ is steeper at lower altitudes than at higher altitudes (Figures 5, 6 and Supplementary Figure S4). In particular, in the two plots at Pitadeniya at 509 m and 618 m, V_sky(θ,α)_ was less than 0.10 in a substantial fraction of the lower one-third of the hemisphere. In contrast, in the higher altitude plots, V_sky(θ,α)_ was greater than 0.10 in a greater fraction of the lower hemisphere. The transition from higher to lower V_sky_, which is a measure of decreasing radiation penetration at lower solar elevations during the morning and evening, was steeper in the higher altitude plots than at lower altitudes (Figure 5 and Supplementary Figure S4). The mid-altitude plots (i.e. Sinharaja-Enasalwatte, Rilagala and Hakgala at 1042 – 1804 m) were intermediate in terms of their canopy openness in the top one-third of the hemisphere and the steepness of its progressive reduction in the middle and lower thirds. Similar variations in canopy openness at different zenith angles have been interpreted by Smith et al. (1992) from contour maps of canopy openness in a lowland moist tropical forest and by Silbernagel and Moeur (2001) in a temperate forest.

**Figure 5:**
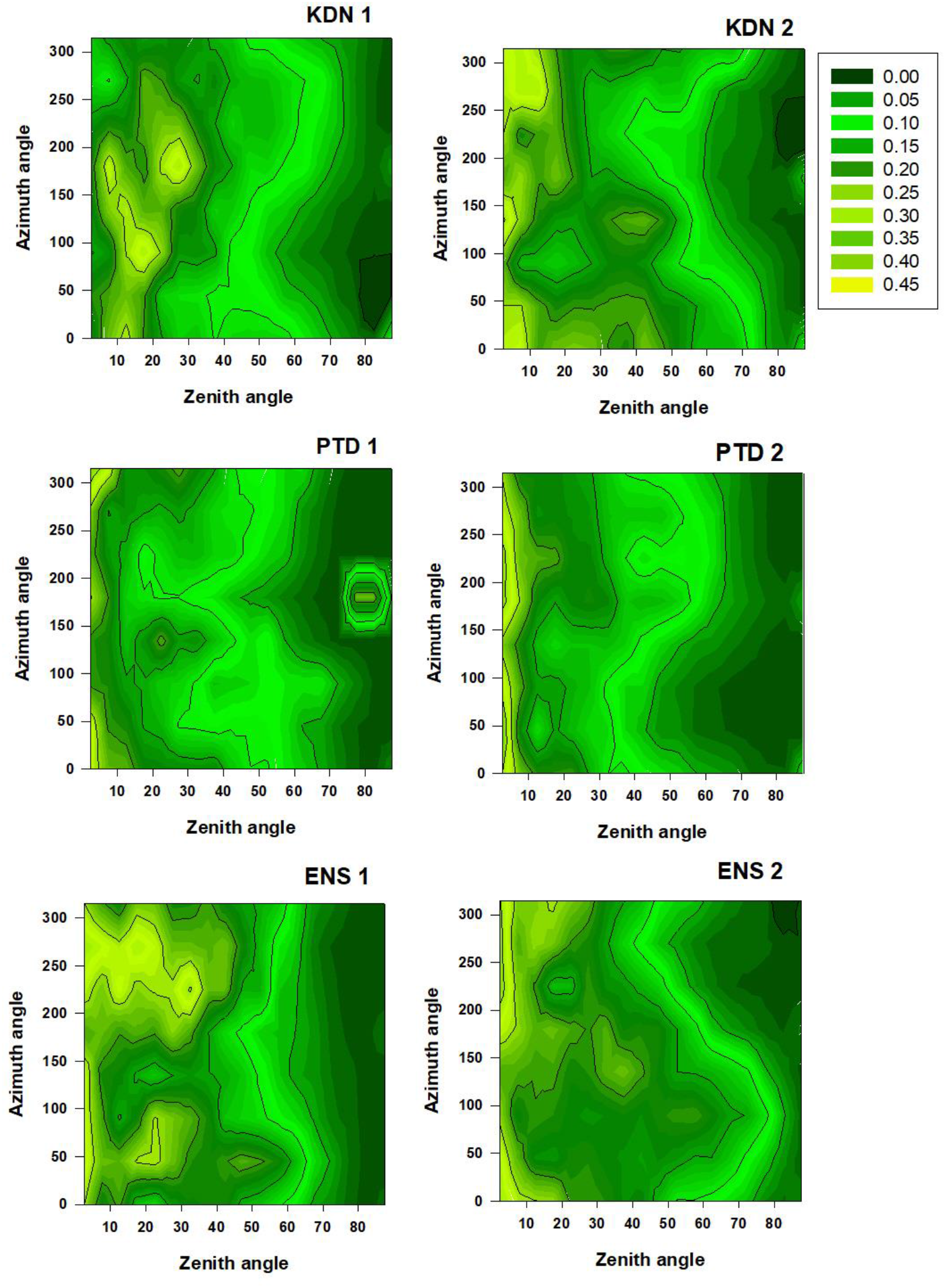
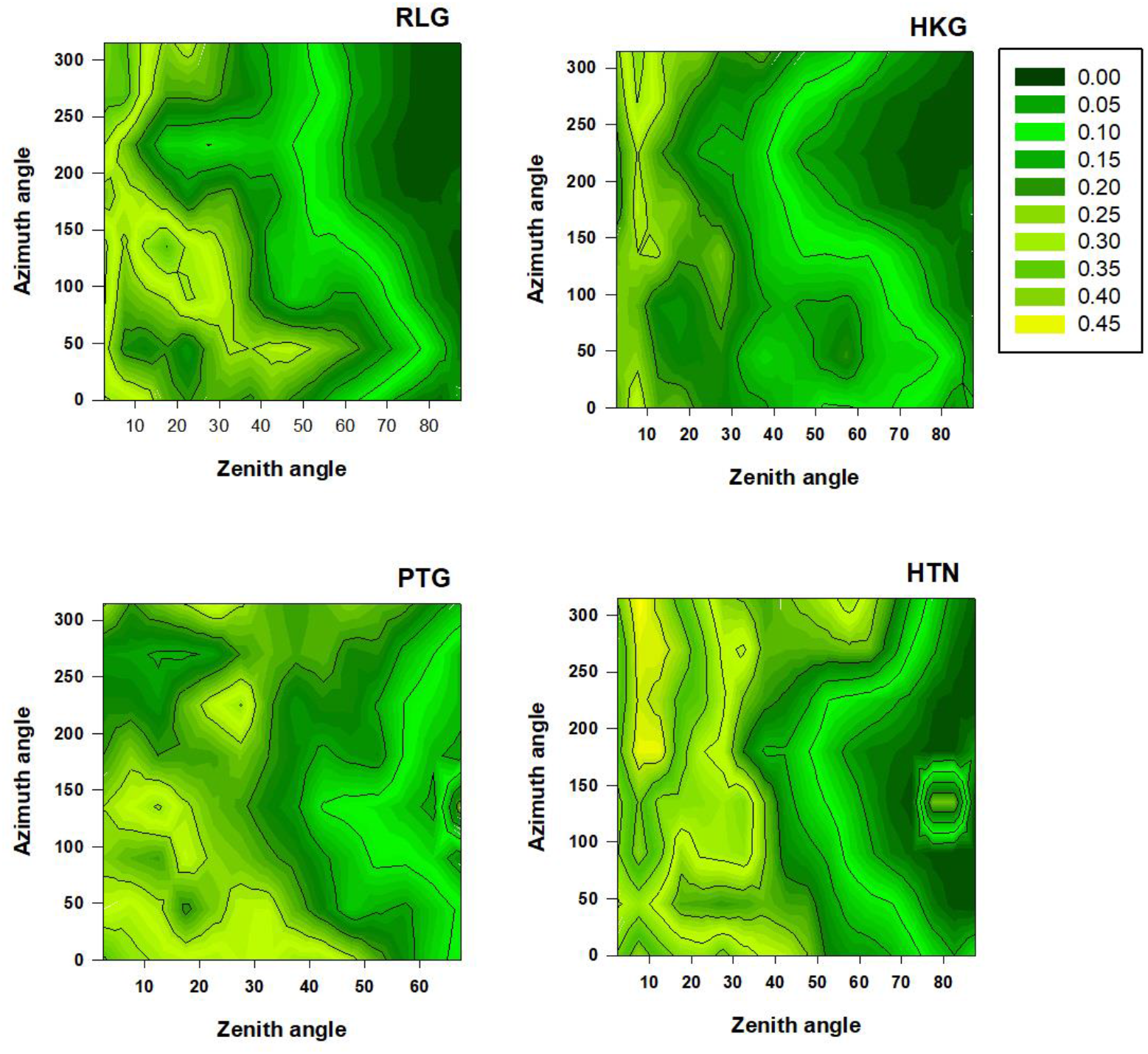
Distribution of the fraction of visible sky (Vsky) across zenith and azimuth angles of the canopy hemisphere in different permanent sampling plots. KDN 1 & 2: Kanneliya; PTD 1 & 2: Sinharaja-Pitadeniya; ENS 1 & 2: Sinharaja-Enasalwatte; RLG: Rilagala; HKG: Hakgala; PTG: Pidurutalagala; HTN: Horton Plains.

The LMM analysis of the overall V_sky(θ,α)_ dataset showed that the random effect of altitude (β) was highly-significant (*p*<0.0001) during all three measurement rounds (γ). The repeated factor, γ, also had a highly-significant (*p*=0.0045) effect on V_sky(θ,α)_. While the γ × θ × α interaction effect was not significant (*p*>0.05), the θ × α interaction effect was significant (*p*=0.0022). The main effects of θ and α also were highly significant (*p*<0.0001). When the LMM analysis was performed for different PSPs, the effect of γ was significant (*p*<0.05) in all PSPs except KDN 2 and HKG (Supplementary Table S1). In all PSPs, the θ × α interaction effect and the main effects of θ and α also were significant.

### Relationship between Variation of Canopy Openness with Azimuth and Aspect of Forest Plots

The specific variation patterns of canopy openness (i.e. V_sky(θ,α)_) with azimuth can be related to the aspect of the forest plots (i.e. the direction of terrain) and the presence or absence of geographical features (e.g. mountains) that could obstruct the incidence of solar radiation on to the forest canopy. For examining this relationship, variation of V_sky(θ,α)_ with azimuth in the top one-third of the hemisphere (zenith angles 0° - 30°, V_sky(0-30)_) provides the best assessment of canopy openness because variation in this segment is least affected by the presence tree stems/trunks (Supplementary Figure S3) (Gonsamo *et al*., 2013).

#### Comparison of canopy openness between the eastern and western sides

In three forest plots (i.e. KDN 2 at 174 m, ENS 1 at 1042 m and ENS 2 at 1065 m) out of the 10 PSPs, V_sky(0-30)_ on the east (azimuth angle = 90°) was significantly (*p*<0.05) lower than that on the west (azimuth angle = 270°) (Figure 6.a). Out of these three plots, KDN 2 is north-west-facing whereas ENS 1 and 2 are east-facing (Table 1). While the KDN 2 plot was on relatively flat terrain (i.e. slope<10%), ENS 1 and ENS 2 were on gentle slopes (i.e. 11%) along the South-North direction (Table 1). In the humid tropical climate that prevails in these plots, it is likely that these plots receive greater solar radiation energy from the east during the pre-noon period than from the west during the post-noon period. Maps of projected shade on these three plots confirm that they are less shaded during the pre-noon period than during the post-noon period (Figure 7). Measurements of diurnal variation of instantaneous incident solar irradiance from an automated weather station established about 100 m away from ENS 1 and 2 also show that irradiance levels during the pre-noon period is greater than those during the post-noon period (Figure 8). Smith *et al*. (1977) also has observed a similar pattern of incident radiation on east- and west-facing slopes of a tropical mountain. Accordingly, it is possible that the lower canopy openness on the east indicates a canopy structure which is designed to maximize radiation capture from the side which receives greater radiation (i.e. the east).

**Figure 6:**
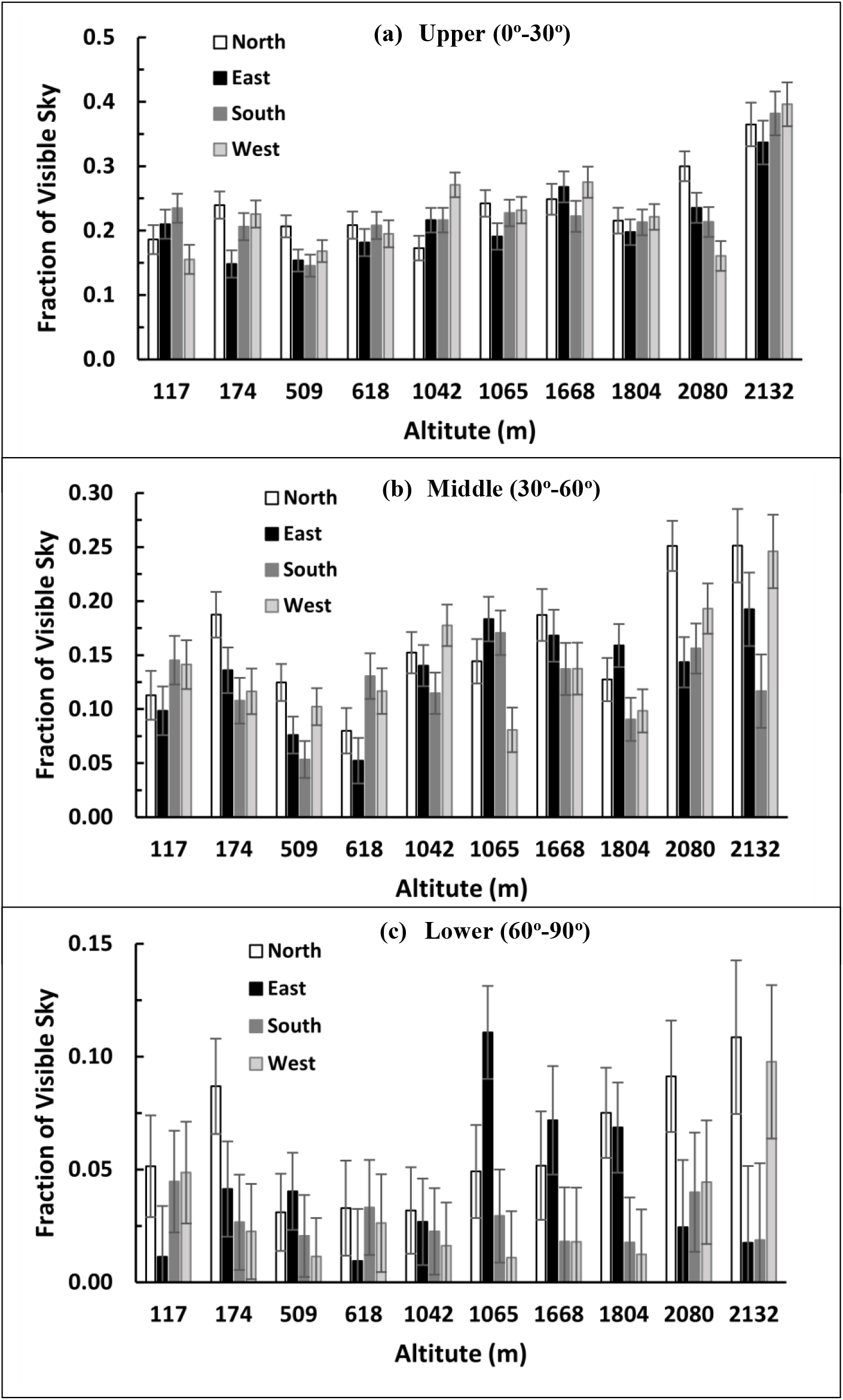
Variation of least square means of visible sky fraction (V_sky_) of forest canopies in four selected directions of the azimuth. Panels a, b and c represent different zenith angle segments of the canopy hemisphere showing V_sky_ at upper, middle and lower segments of the canopy hemisphere. Note different scales of the y-axis in different panels. Error bars are 95% confidence intervals of least square mean.

**Figure 7:**
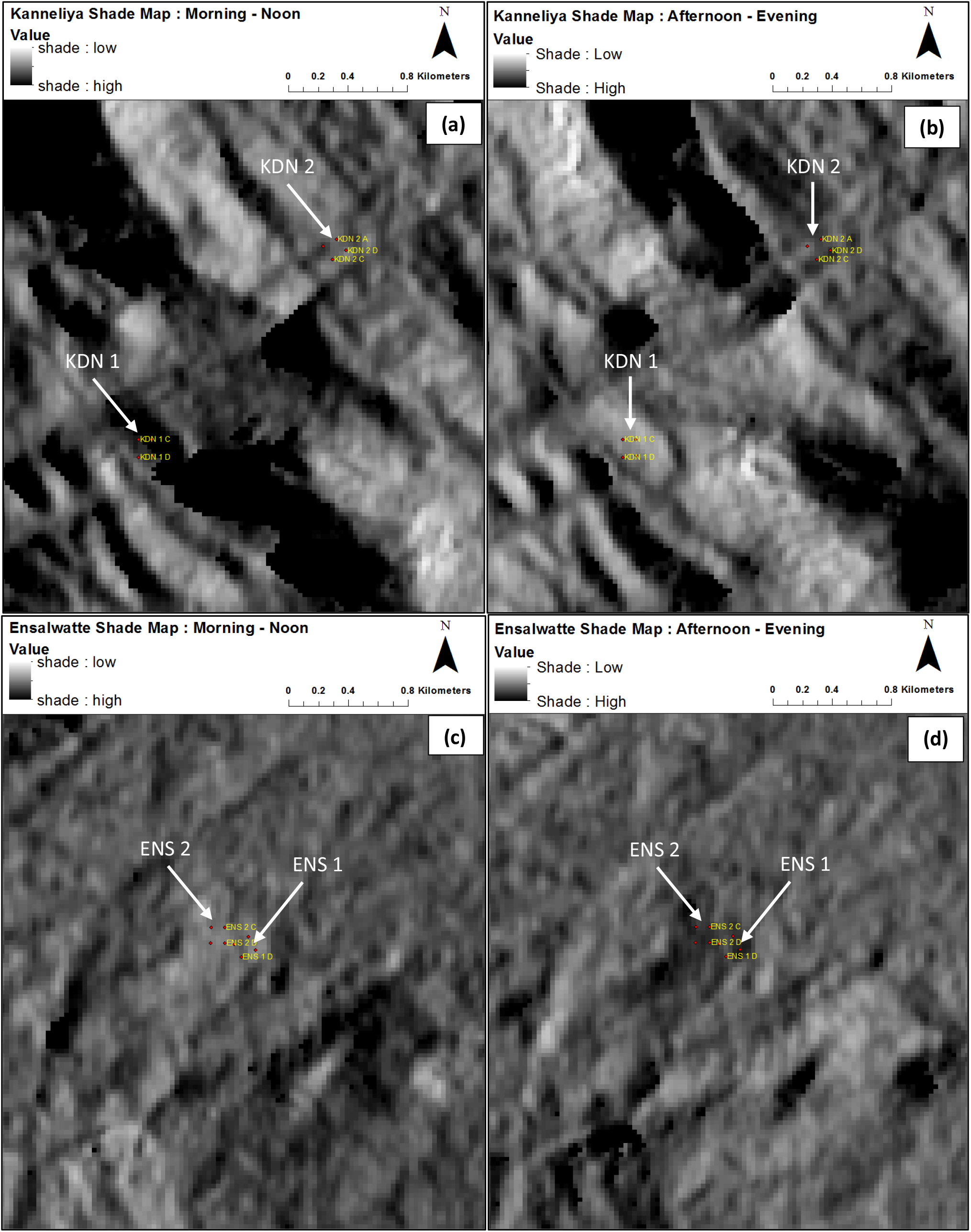
Projected shade on forest canopies in the permanent sampling plots at Kanneliya (KDN) and Sinharaja-Enasalwatte (ENS) forest reserves: (a) Pre-noon KDN; (b) Post-noon KDN; (c) Pre-noon ENS; (d) Post-noon ENS. Shade maps generated with sun angle set at 20° are shown.

**Figure 8:**
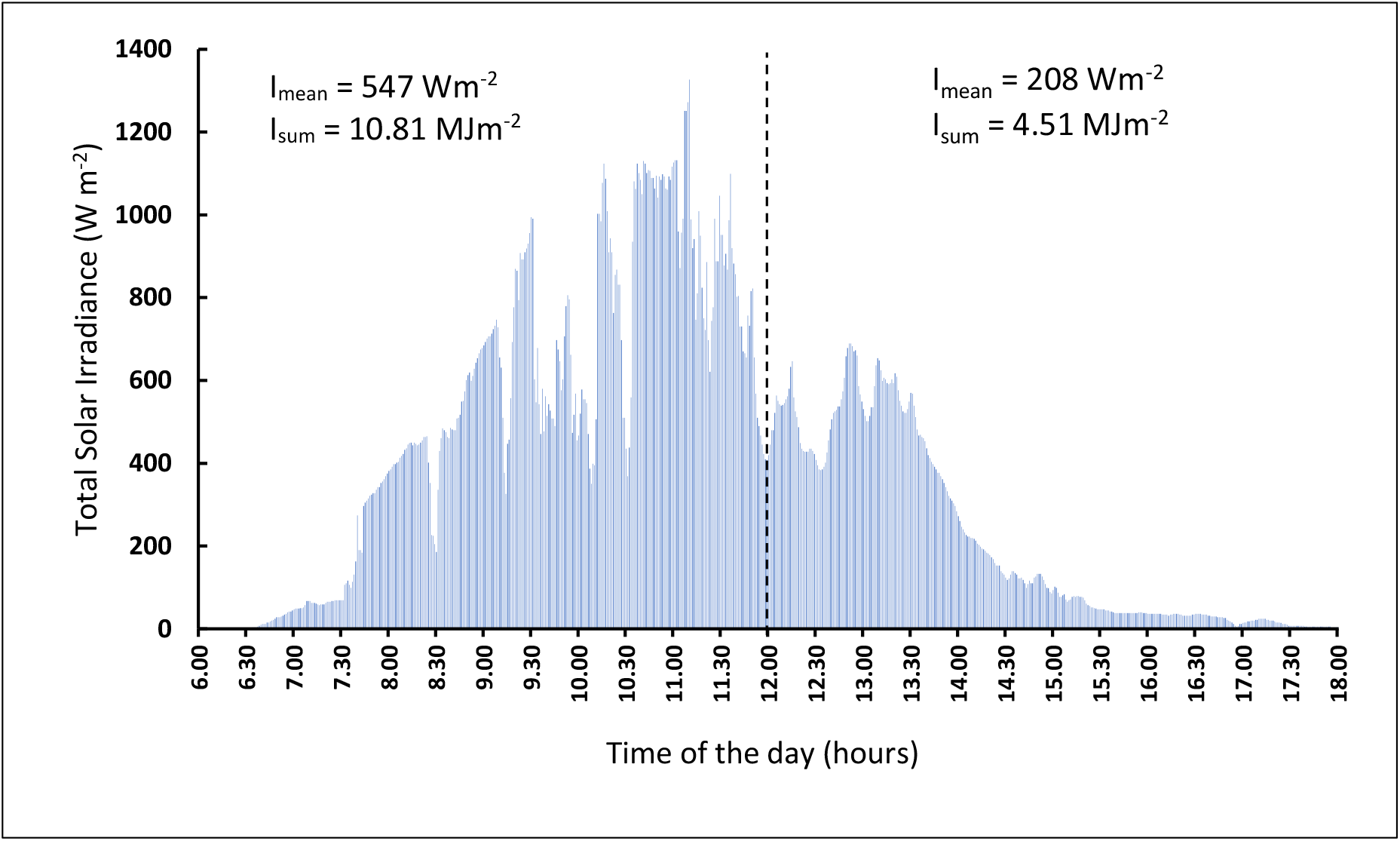
Diurnal variation of total solar irradiance as measured in the automated weather station located near the two permanent sampling plots at Sinharaja-Enasalwatte. Measurements recorded on the 03^rd^ of March, 2022 at one-minute intervals. I_mean_ and I_sum_ are mean irradiance and cumulative total irradiance during pre- and post-noon periods.

The above-described variation pattern observed in KDN 1, ENS 1 and ENS 2 is in agreement with the observations of differential tree growth on east- and west-facing slopes by Liang *et al*. (2006). The east-facing slopes of ENS 1 and 2 (Table 1) probably enhanced this effect. Even though KDN 2 was facing north-west, its very gentle slope was not sufficient to neutralize or reverse the influence of greater irradiance on its east during the M-N period in comparison to the irradiance on its west during the A-E period. Except for KDN 1 and PTG, in which V_sky(0-30)_ on the west was significantly (*p*<0.05) lower than that on the east, the rest of the plots also showed the same pattern shown in KDN 2, ENS 1 and ENS 2, even though the difference was not statistically-significant. In these eight plots, the % increase of V_sky(0-30)_ on the west in comparison to that on the east ranged from 2.7% (RLG) to 52.4% (KDN 2) (Figure 6.a). Except for PTD 2 and HTN, the maps of projected shade on these plots also confirmed the greater irradiance on the east than on the west (Supplementary Figure S5). The fact that PTD 2 and HTN were west-facing slopes (Table 1) could have contributed to the greater morning shade on these plots.

In contrast to the pattern shown in the rest of the PSPs, the forest plots at 117 m at Kanneliya (KDN 1) and at 2080 m at Pidurutalagala (PTG) showed greater V_sky(0-30)_ on the east than on the west (Figure 6.a). This was probably because of the mountains that were present on the eastern sides of these two plots (Table 1). The mountain at PTG was at an aerial distance of 100 m from the plot while at KDN 1 it was 400 m from the plot. These mountains caused shading of the eastern side of the forest canopy during the first half of the day and thus reduced radiation receipt to the east, whereas the western sides received their normal quota during the second half of the day. This was confirmed by the maps of projected shade on these two plots during M-N and A-E periods (Figures 7.a & b and 9.a & b).

**Figure 9:**
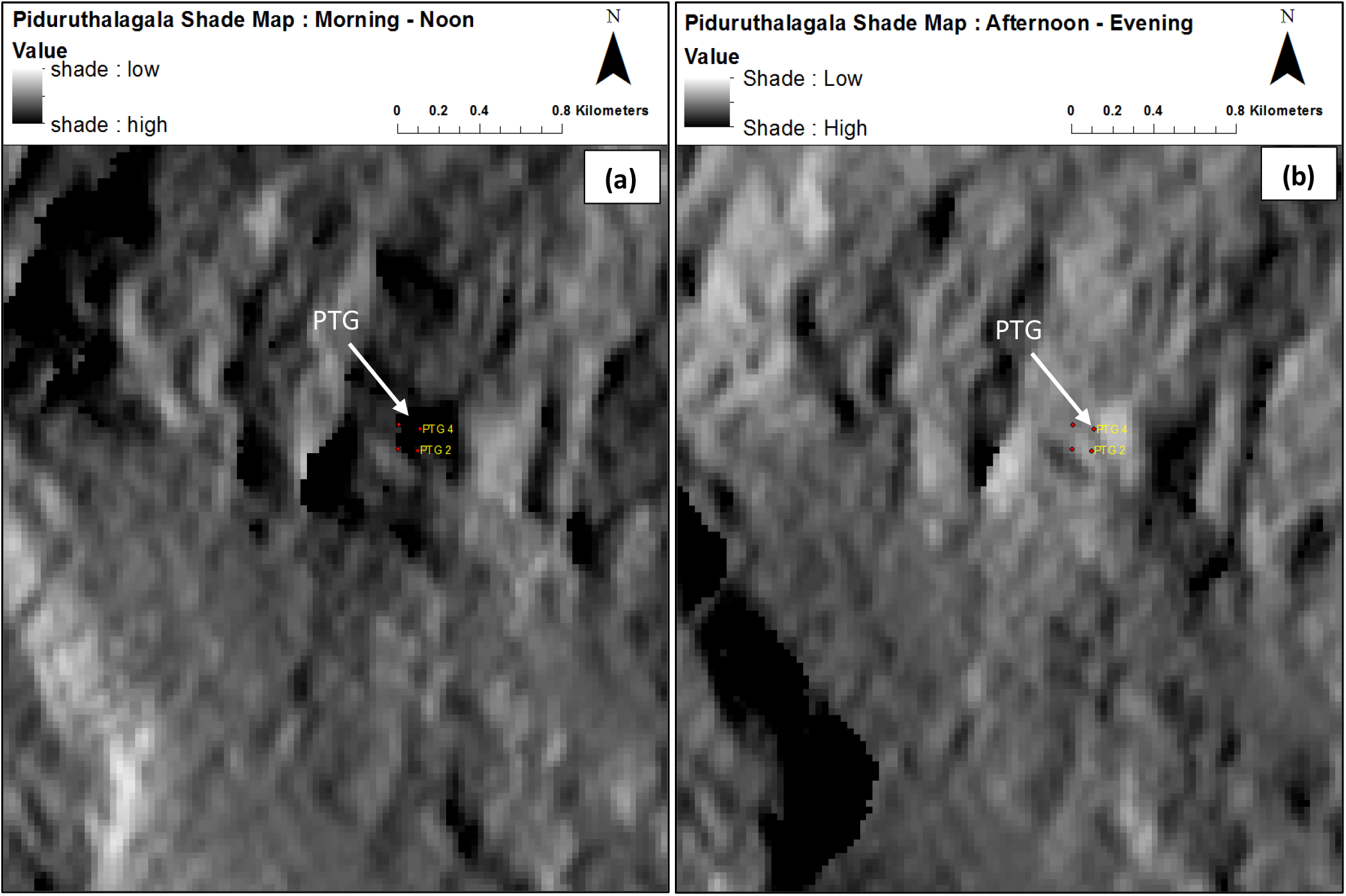
Projected shade on forest canopies in the permanent sampling plot at the Pidurutalagala forest reserve: (a) Pre-noon; (b) Post-noon. Shade maps generated with sun angle set at 20° are shown.

Diurnal variation of irradiance measurements from the automated weather stations established near KDN 1 and PTG also showed greater irradiance during the M-N period than during the A-E period (Figure 10). Accordingly, it is probable that the forest canopies of these two plots allocated a greater proportion of their leaf area to the western side to maximize radiation capture. Consequently, canopy openness was lower on the west than on the east.

**Figure 10:**
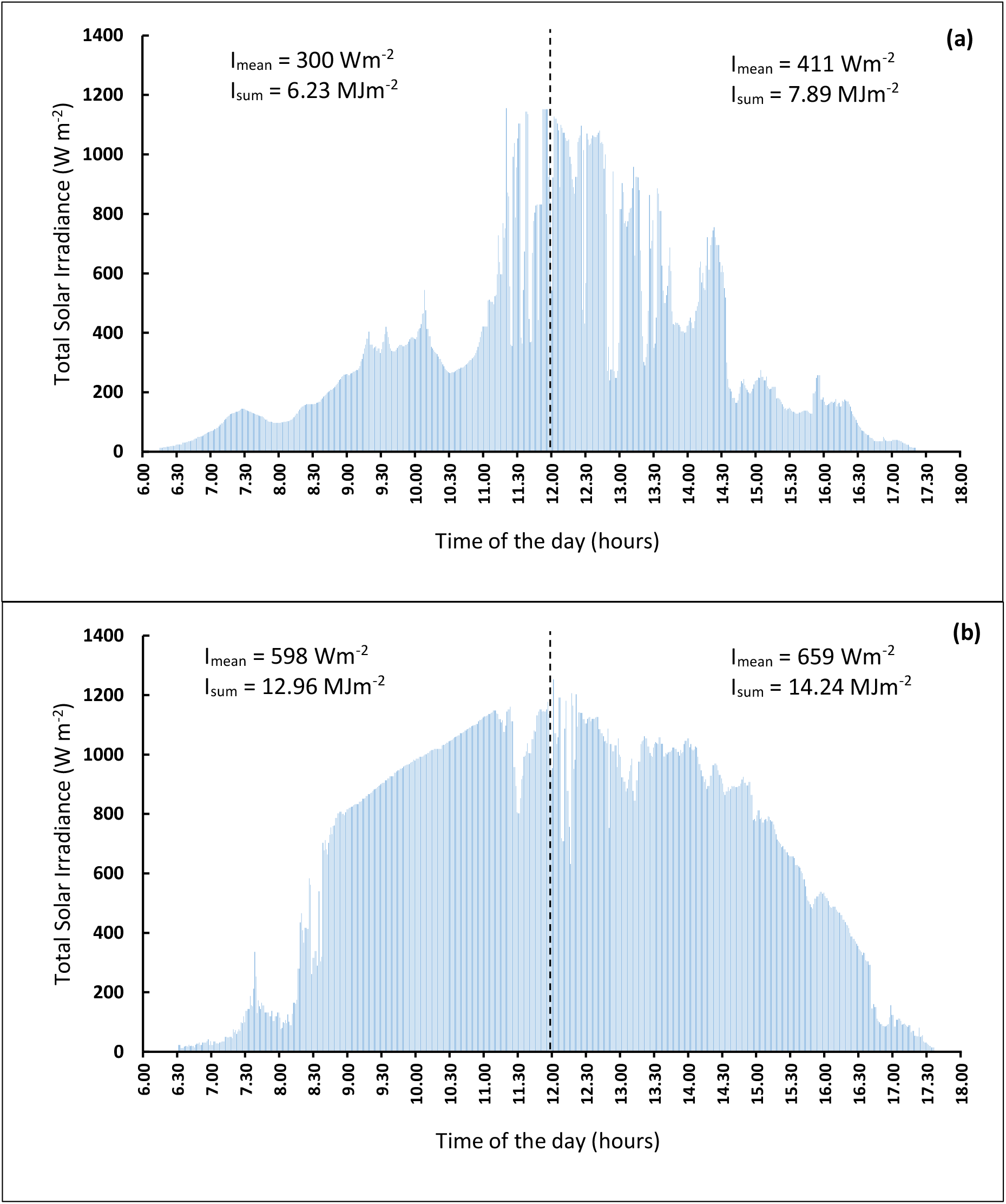
Diurnal variation of total solar irradiance as measured in the automated weather stations located near the permanent sampling plots at Kanneliya Plot 1 (a) and Pidurutalagala (b). Measurements recorded on the 18^th^ of November (Kanneliya) and 18^th^ of October (Pidurutalagala), 2022 at one-minute intervals. I_mean_ and I_sum_ are mean irradiance and cumulative total irradiance during pre- and post-noon periods.

A similar pattern of canopy openness was observed by Frazer *et al*. (2000) in a coastal temperate forest, where the higher canopy openness on the east was attributed to its drier climatic conditions as compared to the west. However, in the humid tropical zone of the present study, wetness of the climate is unlikely to be a determining factor of canopy openness because of the high rainfall regime.

Even though the forest plots in PTD 1 at 509 m, PTD 2 and 618 m, RLG at 1668 m, HKG at 1804 m and HTN at 2132 m showed the expected lower V_sky(0-30)_ on the east in comparison to the west, the difference was not significant (*p*>0.05) (Figure 6.a). While not altering the differential growth of the east- and west-facing sides of the forest canopy in response to differential radiation receipt, effects of local topography could have reduced the difference between the two sides. For example, the two plots at 509 and 618 m at Sinharaja-Pitadeniya, which were on relatively flat terrain had a mountain to the east at a distance of 1 km. The shading effect of this mountain during the first half of the day, while not being as strong as that at Pidurutalagala or Kanneliya, probably reduced the advantage of the eastern side. The forest plot at 1804 m in Hakgala was on the northern slope with a mountain to its south-west. Generally, northern slopes receive less solar radiation intensities than southern slopes because in the northern hemisphere, solar radiation is incident more directly on south-facing slopes (Holland and Steyn, 1975). While its location on the northern slope may not have caused a differential radiation receipt to the eastern and western sides of the forest plot at Hakgala, the mountain to the south-west could have caused a partial shading on the eastern side of the canopy. This is because the azimuth angle of the incident solar radiation during the first half of the day moves from the east to the south so that a mountain to the south would obstruct solar radiation incident on the forest canopy. This shading effect probably reduced the advantage of the eastern side in terms of radiation receipt thus causing the absence of a statistically-significant difference despite showing the expected trend of a lower V_sky(0-30)_ on the east in comparison to the west. The forest plot at Rilagala at 1668 m was on a slightly eastern slope with no obstructing mountains around it. However, the difference in V_sky(0-30)_ between the eastern and western sides was lowest at RLG. This was probably because this is a forest which is at the early stage of recovery from disturbance. A separate study, which involved a detailed vegetation survey (Sanjeewani et al., 2020) showed that around 94% of the tree population in RLG is within the 10 – 20 cm DBH (diameter at breast height) class. Therefore, it is likely that the tree canopy of this forest may not have had adequate time to respond to the differential irradiance on the eastern and western sides.

#### Comparison of canopy openness between the northern and southern sides

Canopy openness in the upper one-third of the hemisphere (V_sky(0-30)_) showed significant (*p*<0.05) variation between the north- and south- facing sides in four out of the 10 PSPs (Figure 6.a). In the plots at 509 m (Sinharaja-Pitadeniya PTD 1) and 2080 m (Pidurutalagala PTG), V_sky(0-30)_ on the south was lower than on the north. This could be a response of the forest canopy to the greater incident radiation on the southern side. Greater allocation of leaf area to the southern side of the canopy would reduce canopy gaps and maximize radiation capture. This is in agreement with the pattern reported by Holland and Steyn (1975). In contrast to these two plots, lower V_sky(0-30)_ on the north than on the south was observed at 117 m (Kanneliya KDN 1) and 1042 m (Sinharaja-Enasalwatte ENS 1). Obstruction of incident radiation by the mountain to the east of the Kanneliya plot (Table 1) probably reduced radiation incidence to the southern part of the forest canopy as well during the first half of the day. This could have shifted the allocation of leaf area from the southern to the northern part of the canopy and consequently a greater V_sky(0-30)_ on the south. The plot at 1042 m (ENS 1) traversed both the southern and northern faces of a hill top. The lower V_sky(0-30)_ on the north probably indicates that the incident radiation was greater on the north-facing portion of the plot which therefore had a greater allocation of leaf area.

In the remaining six plots at 174, 618, 1065, 1668, 1804 and 2132 m, V_sky(0-30)_ was not significantly different between the northern and southern sides of the forest canopy (Figure 6.a). As explained earlier with regard to east versus west differences in V_sky(0-30)_, in these plots, mountains on the east and west (KDN 2 and PTD 2) and south-west (HKG) (Table 1) probably caused reductions in radiation incident on the southern part of the canopy. This caused an even allocation of leaf area on both sides so that V_sky(0-30)_ did not differ significantly. Absence of a significant difference in V_sky(0-30)_ between the northern and southern sides in the plots at 1065 m (Sinharaja-Enasalwatte ENS 2) and 2132 m (Horton Plains HTN) could have been caused by the local climatic variations such as the presence of low-hanging clouds and mist, which would have reduced the radiation incident on the two sides differentially.

#### Canopy openness in the middle and lower thirds of the canopy hemisphere

Estimates of V_sky_ in the middle (zenith angle 30°-60°) and lower (zenith angle 60°-90°) thirds of the canopy hemisphere (Figures 6.b and c) include the effect of tree stems and trunks which get ‘stacked’ in these segments of a hemispherical photograph (Supplementary Figure S3). Therefore, V_sky_ values in these segments are less precise representations of canopy openness than the corresponding values in the upper segment. However, they do represent the degree of canopy openness for radiation incident during the hours of the day which are outside those around the noon (i.e. from sunrise up to *ca.* 1000 hours and from *ca.* 1400 hours until sunset). Accordingly, it is worth examining whether the comparative variation of V_sky_ between the east- and west-facing parts of the forest canopy and that between the northern and southern parts agrees with the expected variation patterns (i.e. greater V_sky_ on the west than east and on the north than south).

Canopy openness in the middle third of the canopy hemisphere (V_sky(30-60)_) was higher on the west than on the east in six out of the 10 PSPs and the difference was statistically-significant (*p*<0.05) in two PSPs (Figure 6.b). Furthermore, V_sky(30-60)_ was greater on the north than on the south in seven PSPs with statistical significance (*p*<0.05) shown in five PSPs. The corresponding variations in canopy openness in the lower third of the hemisphere (V_sky(60-90)_) showed that V_sky(60-90)_ on the east was greater in four PSPs (statistically-significant in one) while that on the north was greater in nine (statistically-significant in four) PSPs (Figure 6.c). While it is possible that variation in stem densities in different PSPs and site-specific topography could have had an influence, the comparative variation patterns of V_sky(30-60)_ and V_sky(60-90)_ provide support to the possibility of forest canopy differentially allocating its leaf area according to the variation of radiation receipt with azimuth.

## CONCLUSIONS

Based on the observations of this work, it is concluded that canopy openness, as quantified by the visible sky fraction of the canopy hemisphere (V_sky_), of tropical rainforests in Sri Lanka increase with increasing altitude, thus indicating a decrease of canopy size, as quantified by the leaf area index. Variation of V_sky_ with azimuth (i.e. angle of direction relative to the compass north) in the series of forest canopies across the altitudinal range from 117 to 2132 m suggests the hypothesis that in a given forest canopy, allocation of leaf area is relatively greater in the directions from which incident radiation is greater. This meant that in situations that do not have obstructions to incoming radiation (e.g. mountains), openness was greater on the: (a) west-facing side of the canopy than on the east-facing side; (b) northern side than on the southern side of the canopy. On the other hand, the presence of obstructions and aspect of the slope of a forest plot could influence the allocation pattern of leaf area and canopy openness of forest canopies to an extent that could reverse the respective patterns observed on relatively flat terrain receiving unobstructed radiation.

## ACKNOWLEDGEMENTS

This work was funded by the National Science Foundation, Sri Lanka (Grant NTRP/2017/CC&ND/TA-04/P-01/01). Assistance in field work from Dineth Dhanushka and Suneth Kanishka is acknowledged.

