## Supplementary_figure_S1 for "Influence of Geographical Aspect and Topography on Canopy Openness in Tropical Rainforests of Sri Lanka along an Altitudinal Gradient"

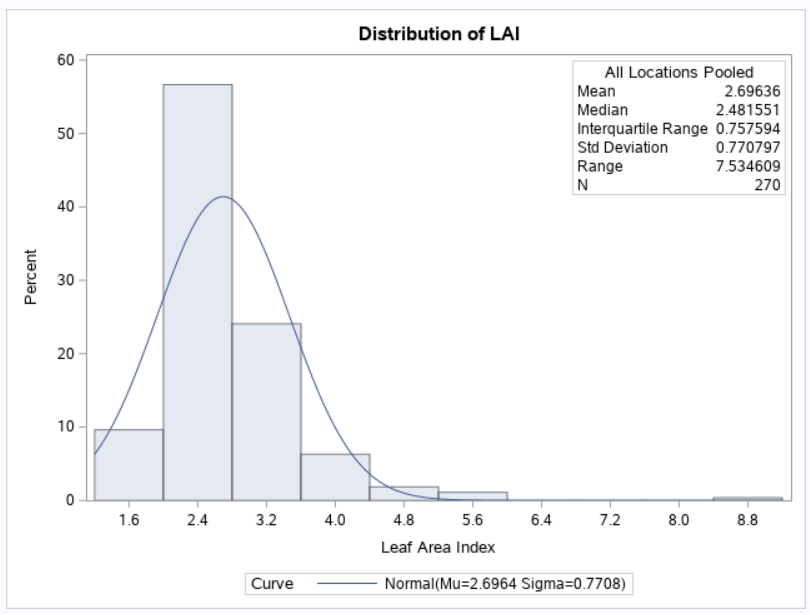

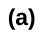

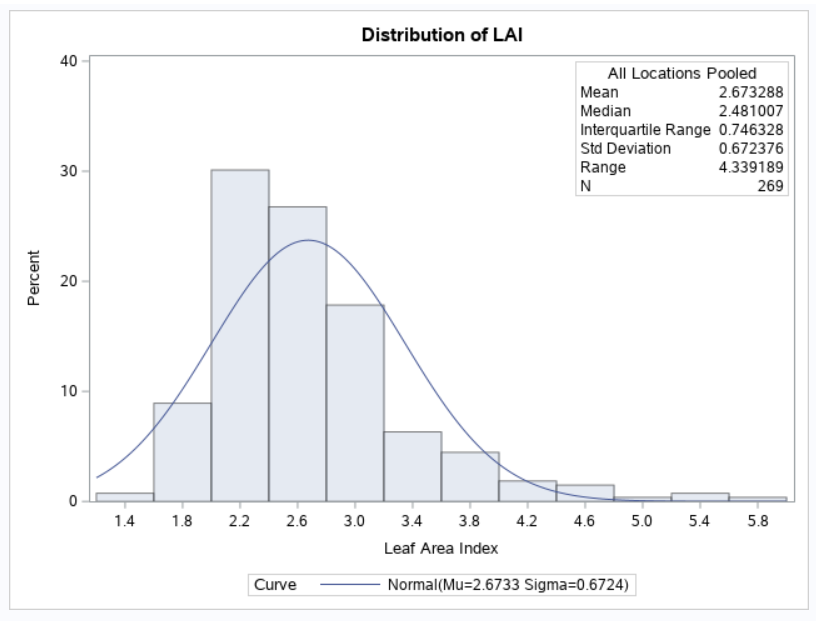

**(b)**

**(C)**

**Supplementary Figure S1:** Distributions of the fraction of visible sky (Vsky) (a) and leaf area index, with (b) and without (c) the outlier

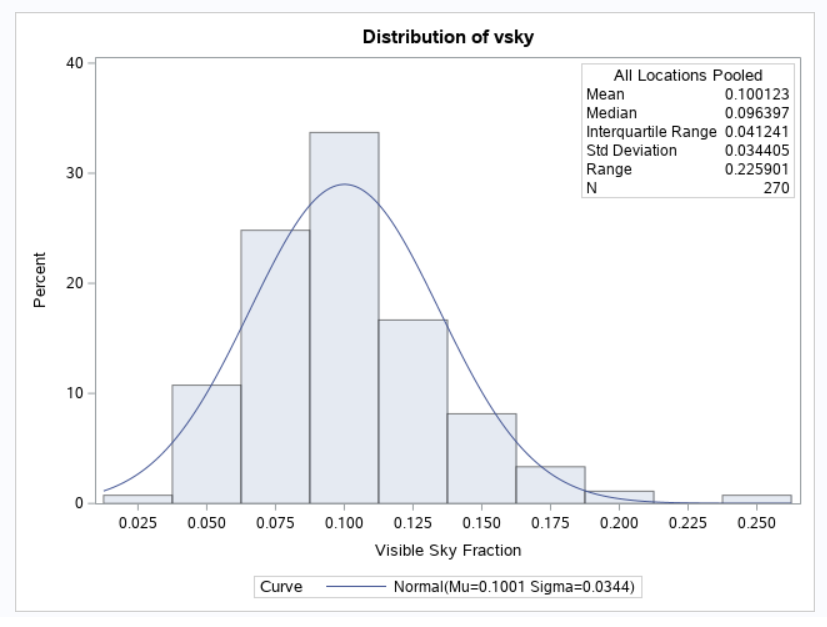

**(a)**
