## Supplementary_figure_S2 for "Influence of Geographical Aspect and Topography on Canopy Openness in Tropical Rainforests of Sri Lanka along an Altitudinal Gradient"

Leaf Area Index

| Altitude (m) | Round 1 | Round 2 | Round 3 |
| --- | --- | --- | --- |
| 117 | 2.74112 | 2.99361 | 3.45885 |
| 174 | 2.5286 | 2.44529 | 2.60172 |
| 509 | 3.11481 | 2.83615 | 3.42279 |
| 618 | 3.38279 | 3.27425 | 3.16188 |
| 1042 | 2.87891 | 2.56908 | 2.43511 |
| 1065 | 2.39605 | 2.33066 | 2.42731 |
| 1668 | 2.24577 | 2.84073 | 2.76707 |
| 1804 | 2.4873 | 2.79012 | 2.94089 |
| 2080 | 1.94011 | 2.43188 | 2.422 |
| 2132 | 2.22446 | 2.14172 | 2.05088 |

| Altitude (m) | Whole |
| --- | --- |
| 117 | 2.74112 |
| 174 | 2.5286 |
| 509 | 3.11481 |
| 618 | 3.38279 |
| 1042 | 2.87891 |
| 1065 | 2.39605 |
| 1668 | 2.24577 |
| 1804 | 2.4873 |
| 2080 | 1.94011 |
| 2132 | 2.22446 |
| 117 | 2.99361 |
| 174 | 2.44529 |
| 509 | 2.83615 |
| 618 | 3.27425 |
| 1042 | 2.56908 |
| 1065 | 2.33066 |
| 1668 | 2.84073 |
| 1804 | 2.79012 |
| 2080 | 2.43188 |
| 2132 | 2.14172 |
| 117 | 3.45885 |
| 174 | 2.60172 |
| 509 | 3.42279 |
| 618 | 3.16188 |
| 1042 | 2.43511 |
| 1065 | 2.42731 |
| 1668 | 2.76707 |
| 1804 | 2.94089 |
| 2080 | 2.422 |
| 2132 | 2.05088 |

**Supplementary Figure S2:** Variation of canopy leaf area index (LAI) with altitude after removing the outlier. Linear and second-order polynomial functions are shown by solid and broken lines respectively.
