## Supplementary_figure_S3 for "Influence of Geographical Aspect and Topography on Canopy Openness in Tropical Rainforests of Sri Lanka along an Altitudinal Gradient"

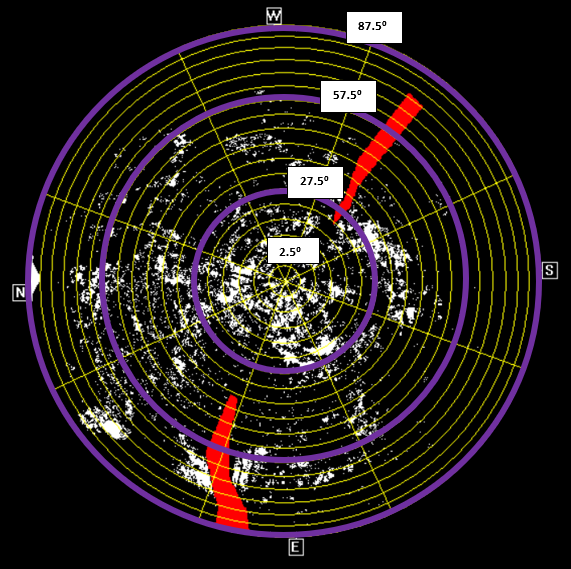

**Supplementary Figure S3:** A canopy hemispherical photograph. The concentric rings denote different zenith angles with the purple-coloured rings showing the three zenith angle segments (0^o^–30^o^, 30^o^–60^o^, 60^o^–90^o^). The radial lines demarcate azimuthal segments with the four main aspects (North, East, South and West) are shown as N, E, S and W respectively. Excluded tree trunks are shown in red.
