## Supplementary_figure_s4 for "Influence of Geographical Aspect and Topography on Canopy Openness in Tropical Rainforests of Sri Lanka along an Altitudinal Gradient"

**Supplementary Figure S4:** Variation of least square means of visible sky fraction (V_sky_) of forest canopies in three zenith angle segments (0^o^–30^o^, 30^o^–60^o^, 60^o^–90^o^) of the canopy hemisphere. Panels a, b, c and d represent the four major aspects of the canopy hemisphere at azimuth angles 0^o^, 90^o^, 180^o^ and 270^o^ respectively. Error bars are 95% confidence intervals of least square mean.

**(a) North**

Fraction of Visible Sky

| Altitude (m) | 0-30 | 30-60 | 60-90 |
| --- | --- | --- | --- |
| 117 | 0.1861 | 0.1128 | 0.05144 |
| 174 | 0.2397 | 0.1874 | 0.08683 |
| 509 | 0.2068 | 0.1247 | 0.031 |
| 618 | 0.2086 | 0.08 | 0.03289 |
| 1042 | 0.1729 | 0.1523 | 0.03183 |
| 1065 | 0.2423 | 0.1443 | 0.04911 |
| 1668 | 0.2486 | 0.1872 | 0.05172 |
| 1804 | 0.2156 | 0.1274 | 0.07511 |
| 2080 | 0.3 | 0.2511 | 0.09128 |
| 2132 | 0.3649 | 0.2513 | 0.1086 |

Fraction of Visible Sky

| Altitude (m) | 0-30 | 30-60 | 60-90 |
| --- | --- | --- | --- |
| 117 | 0.2101 | 0.0985 | 0.01128 |
| 174 | 0.1482 | 0.136 | 0.04133 |
| 509 | 0.1536 | 0.076 | 0.04033 |
| 618 | 0.1815 | 0.05233 | 0.009401 |
| 1042 | 0.2162 | 0.1403 | 0.02678 |
| 1065 | 0.191 | 0.1834 | 0.1107 |
| 1668 | 0.2679 | 0.168 | 0.07178 |
| 1804 | 0.1976 | 0.1589 | 0.06856 |
| 2080 | 0.2354 | 0.1434 | 0.02439 |
| 2132 | 0.3368 | 0.1924 | 0.01756 |

Fraction of Visible Sky

| Altitude (m) | 0-30 | 30-60 | 60-90 |
| --- | --- | --- | --- |
| 117 | 0.2347 | 0.1453 | 0.04461 |
| 174 | 0.2062 | 0.1078 | 0.02661 |
| 509 | 0.1456 | 0.05344 | 0.02053 |
| 618 | 0.2081 | 0.1306 | 0.03317 |
| 1042 | 0.2163 | 0.1147 | 0.02256 |
| 1065 | 0.2275 | 0.1707 | 0.02939 |
| 1668 | 0.2222 | 0.1372 | 0.01806 |
| 1804 | 0.2129 | 0.09056 | 0.01761 |
| 2080 | 0.2134 | 0.1561 | 0.03995 |
| 2132 | 0.3821 | 0.1167 | 0.01878 |

**(b) East**

**(c) South**

Fraction of Visible Sky

| Altitude (m) | 0-30 | 30-60 | 60-90 |
| --- | --- | --- | --- |
| 117 | 0.1553 | 0.1412 | 0.04861 |
| 174 | 0.2258 | 0.1165 | 0.0225 |
| 509 | 0.1682 | 0.1023 | 0.01139 |
| 618 | 0.1951 | 0.1167 | 0.02622 |
| 1042 | 0.2711 | 0.1776 | 0.01622 |
| 1065 | 0.2318 | 0.08083 | 0.01089 |
| 1668 | 0.2752 | 0.1375 | 0.01794 |
| 1804 | 0.2213 | 0.09844 | 0.01233 |
| 2080 | 0.1607 | 0.1931 | 0.04436 |
| 2132 | 0.3962 | 0.246 | 0.09767 |

**(d) West**
