## Supplementary_figure_S5 & S6 for "Influence of Geographical Aspect and Topography on Canopy Openness in Tropical Rainforests of Sri Lanka along an Altitudinal Gradient"

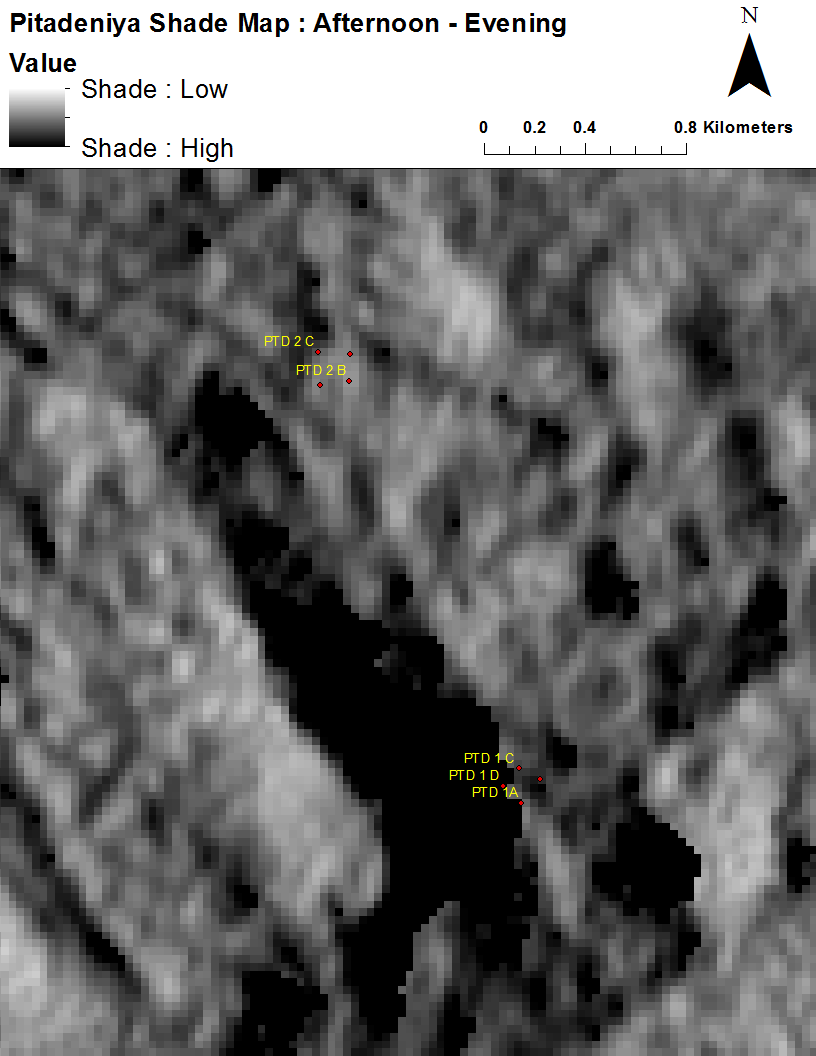

**(b)**

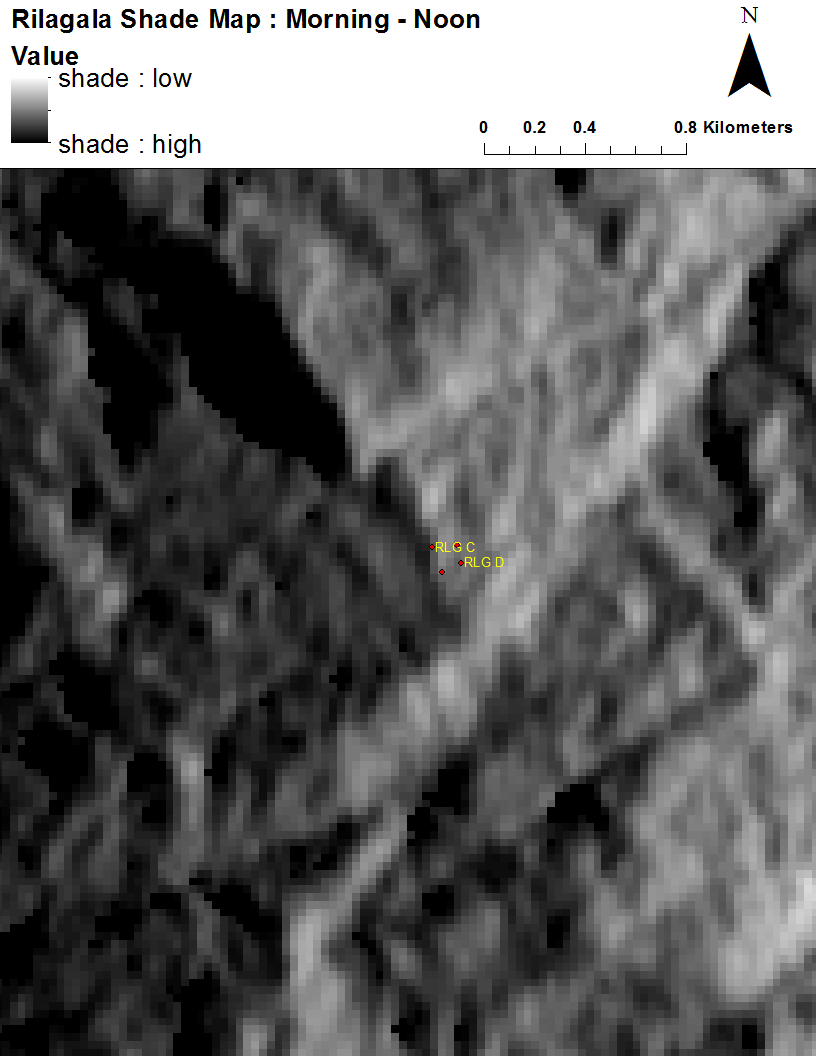

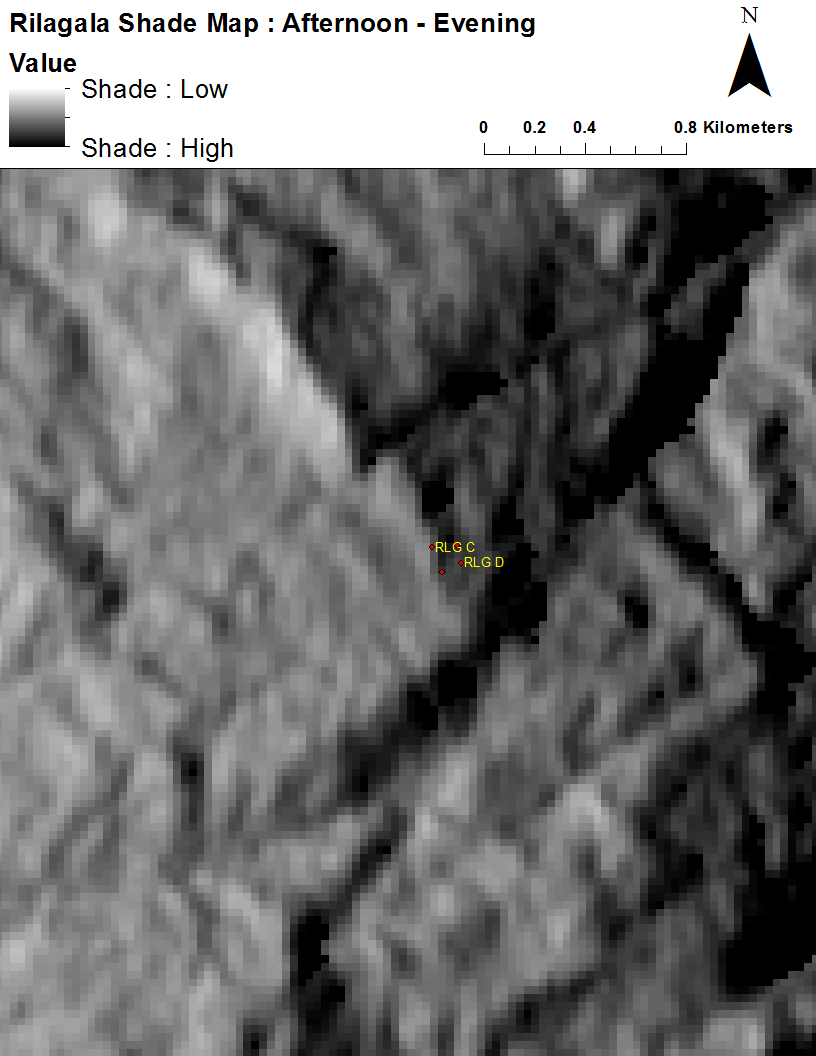

**(c)**

**(d)**

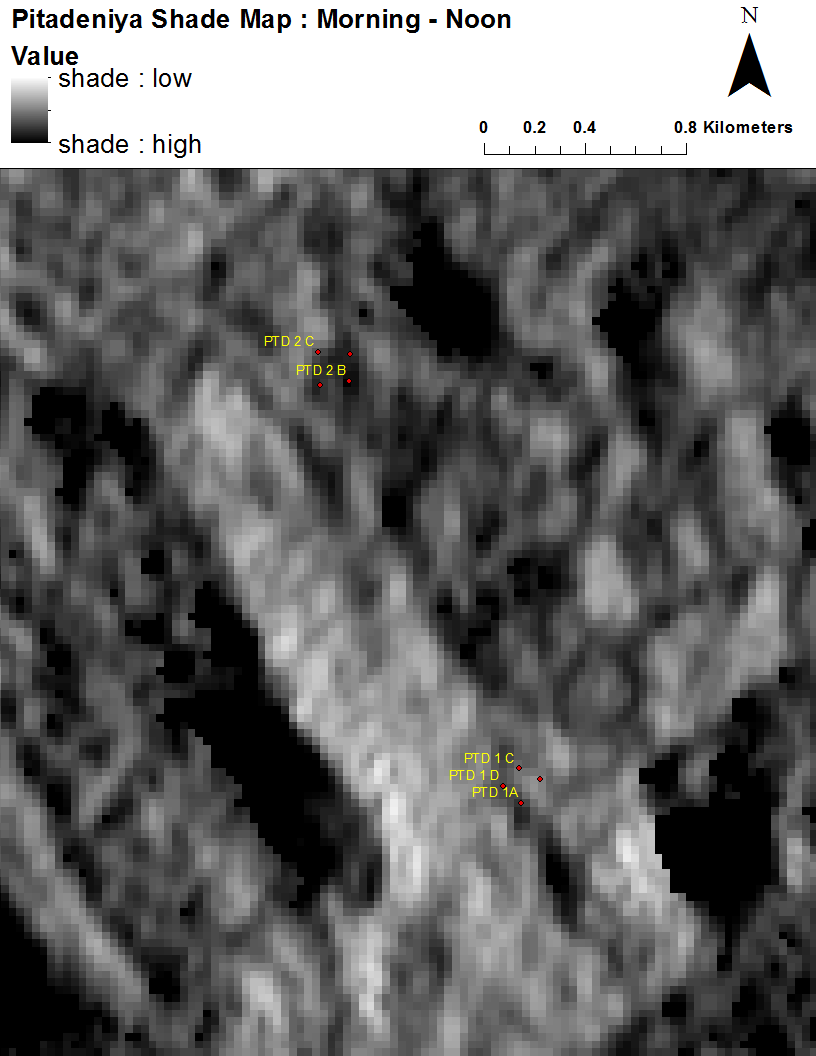

PTD 2

**(a)**

PTD 2

PTD 1

PTD 1

RLG

RLG

**Supplementary Figure S5:** Projected shade on forest canopies in the permanent sampling plots in the forest reserves at Sinharaja-Pitadeniya (PTD) and Rilagala (RLG): (a) Pre-noon PTD; (b) Post-noon PTD; (c) Pre-noon RLG; (d) Post-noon RLG. Shade maps generated with sun angle set at 20^o^ are shown.

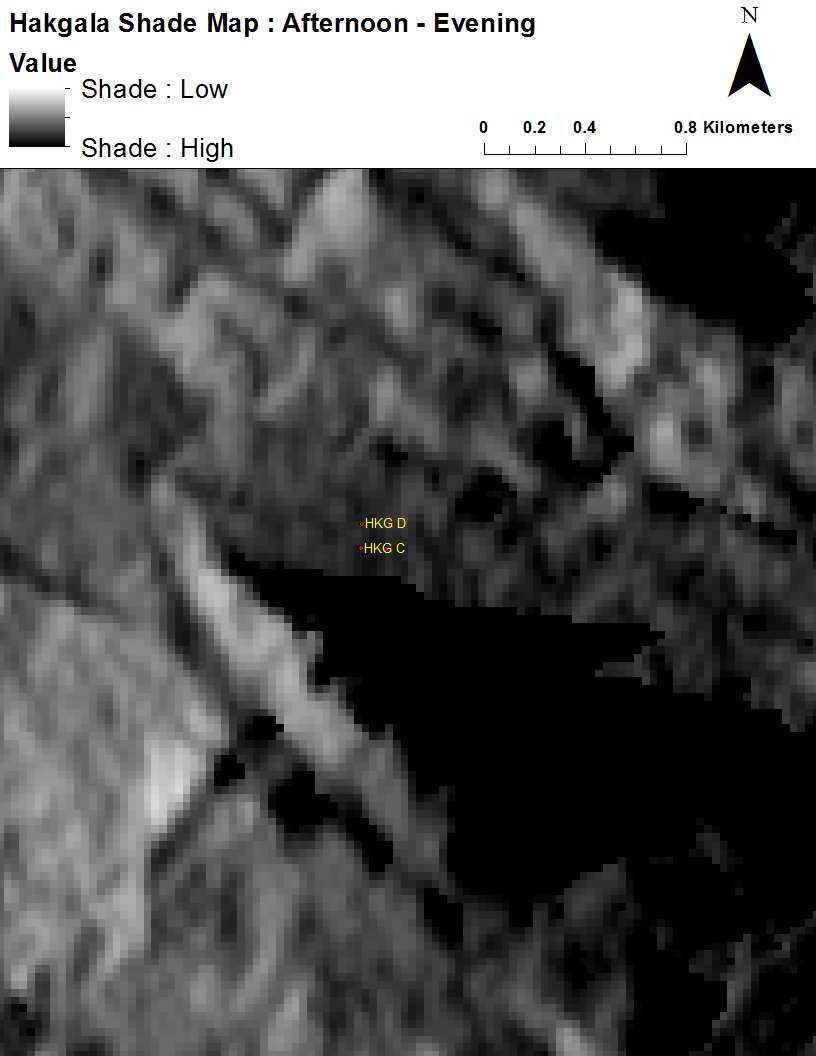

**(f)**

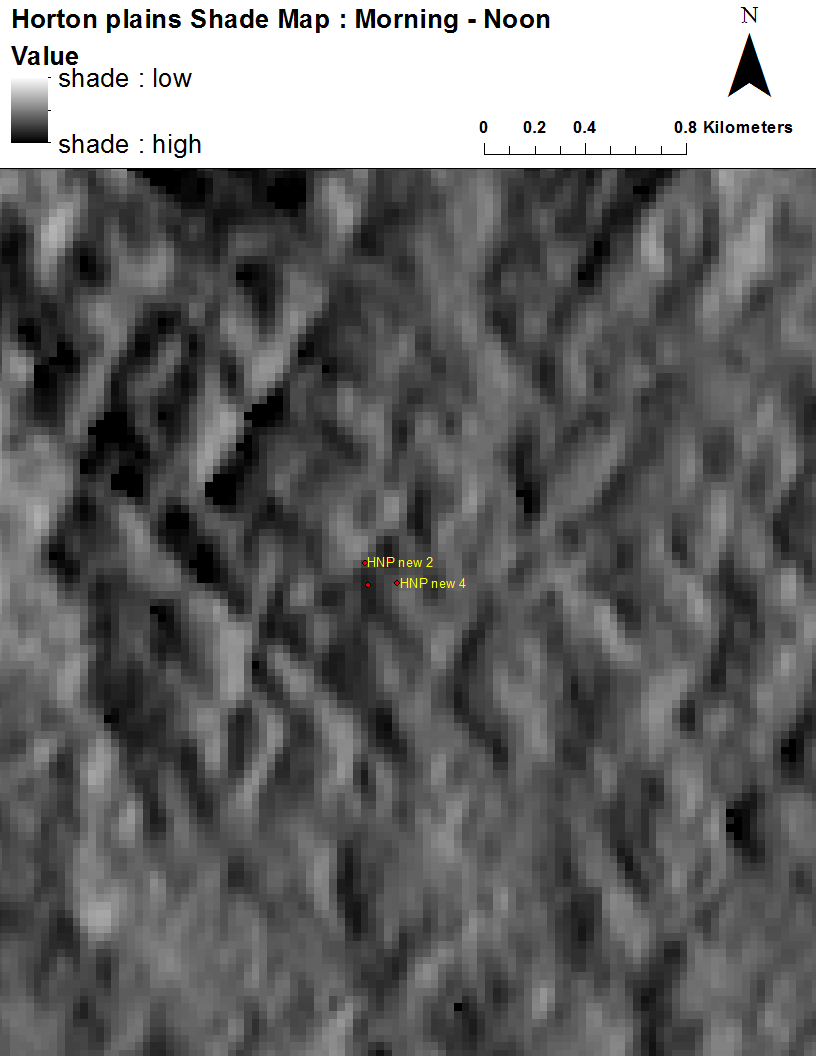

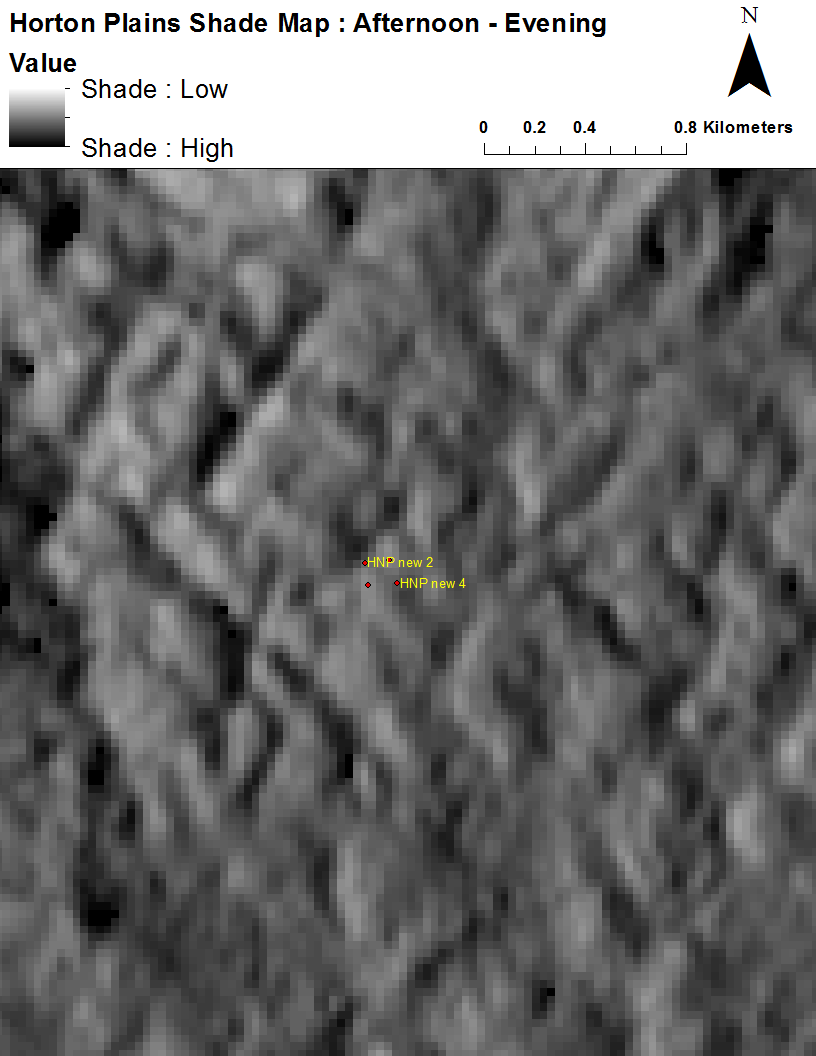

**(g)**

**(h)**

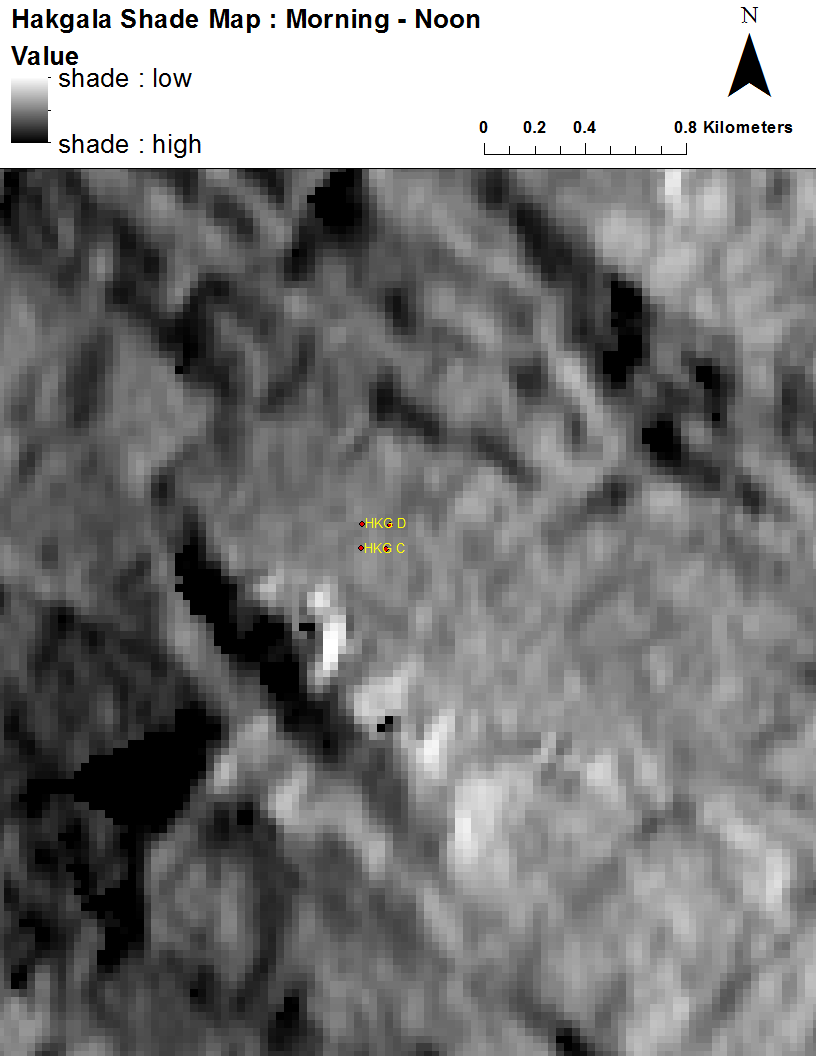

**(e)**

HKG

HKG

HTN

HTN

**Supplementary Figure S5 *continued*:** Projected shade on forest canopies in the permanent sampling plots in the forest reserves at Hakgala (HKG) and the Horton Plains National Park (HNP): (e) Pre-noon HKG; (f) Post-noon HKG; (g) Pre-noon HTN; (h) Post-noon HTN. Shade maps generated with sun angle set at 20^o^ are shown.
