## Supplementary_table_S1 for "Influence of Geographical Aspect and Topography on Canopy Openness in Tropical Rainforests of Sri Lanka along an Altitudinal Gradient"

**Supplementary Table S1.** Significance of the fixed effects of measurement round, zenith angle segment and azimuth angle on visible sky fraction (V_sky_) of canopy hemispherical segments of tropical rainforests of Sri Lanka at different altitudes

| Altitude (m) | ^†^Significance levels of fixed effects of V_sky(θ,α)_ | | | |
| --- | --- | --- | --- | --- |
|  | Round (γ) | Zenith angle segment (θ) | Azimuth angle (α) | θ x α |
| 117 | <0.0001 | <0.0001 | 0.0018 | <0.0001 |
| 174 | ns | <0.0001 | <0.0001 | <0.0001 |
| 509 | 0.0483 | <0.0001 | <0.0001 | 0.0003 |
| 618 | <0.0001 | <0.0001 | <0.0001 | 0.0272 |
| 1042 | 0.0034 | <0.0001 | <0.0001 | <0.0001 |
| 1065 | 0.0003 | <0.0001 | <0.0001 | <0.0001 |
| 1668 | 0.0143 | <0.0001 | <0.0001 | 0.0287 |
| 1804 | ns | <0.0001 | <0.0001 | <0.0001 |
| 2080 | <0.0001 | <0.0001 | <0.0001 | <0.0001 |
| 2132 | 0.0104 | <0.0001 | <0.0001 | 0.0006 |

^†^Estimated based on a linear mixed model analysis.
